# Same data, different results? Machine learning approaches in bioacoustics

**DOI:** 10.1101/2024.04.14.589403

**Authors:** Kaja Wierucka, Derek Murphy, Stuart K Watson, Nikola Falk, Claudia Fichtel, Julian León, Stephan T Leu, Peter M Kappeler, Elodie F Briefer, Marta B Manser, Nikhil Phaniraj, Marina Scheumann, Judith M Burkart

## Abstract

Automated acoustic analysis is increasingly used in behavioural ecology, and determining caller identity is a key element for many investigations. However, variability in feature extraction and classification methods limits the comparability of results across species and studies, constraining conclusions we can draw about the ecology and evolution of the groups under study. We investigated the impact of using different feature extraction (spectro-temporal measurements, linear and Mel-frequency cepstral coefficients, as well as highly comparative time-series analysis) and classification methods (discriminant function analysis, neural networks, random forests, and support vector machines) on the consistency of caller identity classification accuracy across 16 mammalian datasets. We found that Mel-frequency cepstral coefficients and random forests yield consistently reliable results across datasets, facilitating a standardised approach across species that generates directly comparable data. These findings remained consistent across vocalisation sample sizes and number of individuals considered. We offer guidelines for processing and analysing mammalian vocalisations, fostering greater comparability, and advancing our understanding of the evolutionary significance of acoustic communication in diverse mammalian species.

## INTRODUCTION

Many animals produce individually distinct calls – vocalisations that contain unique characteristics or features, allowing for differentiation and identification of individuals (Yorzinski, 2017). Conveying information about individual identity is widespread among taxa, and particularly prevalent in social species, as identity plays a pivotal role in regulating various social interactions (Tibbetts & Dale, 2007; Wyman et al., 2022). The ability for researchers to accurately and reliably identify individuals based on their acoustic cues is also immensely important, serving as a gateway to a myriad of applications, e.g., to gain insights into the behaviour and ecology of species or to inform conservation and management strategies (Gentry et al., 2020; Terry et al., 2005; Tuia et al., 2022).

Quantifying the degree of individual identity information that is encoded in a particular vocalisation is difficult. Broadly, the process requires recording sound (multiple vocalisations per individual on separate occasions), extracting and quantifying their acoustic features (‘feature extraction’) from the recording and using these features to classify the vocalisation as belonging to one of a number of possible individuals (Linhart et al., 2022). Recently, with greater computational power becoming more and more accessible, new, automated methods for feature extraction and classification are becoming increasingly popular (Mcloughlin et al., 2019; Mutanu et al., 2022; Rutz et al., 2023). The ease of application of various classifiers, including those using machine learning techniques, has provided many opportunities within the field of bioacoustics (Phaniraj et al., 2023; Stowell et al., 2019; Valletta et al., 2017). As a consequence, a plethora of methods is currently used with no consensus or consistent protocol (Linhart et al., 2019; Odom et al., 2021) (although, some general guidelines and good practices have been put forward (Linhart et al., 2022; Odom et al., 2021; Oswald et al., 2022)). One reason could be that individuality can be encoded in different ways, which varies by species and context, making it difficult to generalise methods (Stowell et al., 2019). Furthermore, the set of methods that can be used is influenced by various limitations of each particular dataset (e.g., sample size, acoustic characteristics of given vocalisation) vary depending on the research question being investigated. Consequently, most researchers need to develop their own pipelines, which leads to various, non-compatible and often unjustified methods for investigating individual identity (Linhart et al., 2022). This makes it nearly impossible make comparisons across studies or expand conclusions beyond one study system.

Prior studies have explored the efficiency of various methods for determining individual identity information, yet have mostly focused on finding the best method for a given dataset, context or species (Clink et al., 2020; Clink & Klinck, 2021; Pérez-Espinosa et al., 2018; Phaniraj et al., 2023; Zhang et al., 2018). While these studies report important findings, there is no information about the potential variation in results that may derive from using different extraction and classification method combinations for determining individual identity in bioacoustics. Previous research has drawn attention to this issue and call for a unification of the methods used (Keen et al., 2021; Oswald et al., 2022) and several excellent studies have investigated the impact of different statistical (Linhart et al., 2019, 2022) or feature extraction methods (Clemins et al., 2005) on the degree of individual identity quantified in vocalizations (Clemins et al., 2005). However, none of the above have considered the combined effects of feature extraction and classification methods on results. A comparison of the consistency of results obtained using various extraction and classification methods, and a validation of the relationship between these would provide a first step towards the unification of methodologies on vocal individuality, thus ensuring robust and reliable, comparable inferences about the data and, by extension, animal communication, and connected fields.

Research on mammalian vocalisations remains relatively limited compared to the well-established field of avian bioacoustics. To address this gap, in this paper, we begin by systematically reviewing the prevailing methods employed for identifying individual vocal distinctiveness in mammals, aiming to discern the diversity of approaches. Subsequently, using data spanning a diverse range of mammalian taxa, we undertook a broadscale analysis to assess the relative differences in results obtained using various combinations of feature extraction and classification methods for the classification of vocalisations to individual identity in mammals. Additionally, we examined the influence of sample size (number of vocalisations per individual) and number of classes (number of individuals to classify) on these outcomes. The main aim was to establish which methods consistently produce the least amount of variation in results, in order to identify the feature extraction and classification methods that produce robust results. Finally, we present guidelines for future studies, with the goal of fostering a more unified approach to the investigation of animal bioacoustics.

## MATERIALS AND METHODS

### LITERATURE REVIEW

To obtain detailed information about the frequency of use and diversity of feature extraction and classification methods used for quantifying individual identity in mammalian vocalisations, we conducted a systematic literature review. We extracted the first 1000 publications in Google Scholar using the search term “Individual acoustic OR vocal”, limiting the search to articles published in years 2000-2022 (search conducted on the 18^th^ of August 2022). From this list, in our analysis, we included only empirical studies (reviews were excluded) that included an analysis of acoustic cues and an assessment of whether they are individually distinct and targeted a mammalian species. If multiple methods were used in the same study, we treated each method/combination of methods, and its results as a separate entry. In instances where studies provided both cross-validated and non-cross-validated values, we exclusively utilised the cross-validated data for our analysis.

Our examination centred on the extraction methods, classifiers, and the targeted species. With a foundation in the source-filter theory of mammalian acoustics, we scrutinized studies for their employed feature extraction methods, identifying whether they were source-(spectral), or filter-related (features describing the spectral envelope such as formants or MFCC), or a combination of both. Additionally, we catalogued the specific acoustic parameters used, assessing the justifications provided for their selection, which fell into three categories: internal (detailed feature descriptions and rationale), external (citations from other works), or none. Furthermore, we delved into the classifiers employed across the studies listing all possible methods used. We also scrutinised whether the assumptions of the most common classifier were appropriately checked and met. Finally, our review extended to the diversity of taxa, examining the targeted orders and genera among the articles analysed.

### COMPARISON OF FEATURE EXTRACTION AND CLASSIFICATION METHODS

#### Data collection

To make our findings as widely applicable as possible, we used data from 14 mammalian species: alpaca (*Vicugna pacos*), domestic cat (*Felis catus*), Egyptian fruit bat (*Rousettus aegyptiacus*), domestic goat (*Capra aegagrus hircus*), domestic horse (*Equus caballus*), sooty mangabey (*Cercocebus atys*), common marmoset (*Callithrix jacchus*), meerkat (*Suricata suricatta*), grey mouse lemur (*Microcebus murinus*), domestic pig (*Sus scrofa domesticus*), redfronted lemur (*Eulemur rufifrons*), domestic sheep (*Ovis aries*), squirrel monkeys (*Saimiri sciureus*) and tree shrew (*Tupaia belangeri*) (for detailed information on each dataset, please refer to the Data Sources section). For two species (marmosets and mouse lemurs), we had recordings of two distinct call types, and these were treated as independent datasets. Consequently, our analysis incorporated a total of 16 unique datasets, which we refer to as ‘species’ in the model descriptions. For each species, data were limited to the same call type (all were of relatively constant frequency) and each file contained a single call (call bouts were not included in the analysis), with the recording cropped tightly around the vocalisation. Caller identity was known for all calls. Datasets varied in the number of calls per individual and the number of individuals they included, thus, we prepared two separate datasets for the different analyses we conducted. As a large number of recordings per individual is often difficult to obtain, and we wanted to include a wider range of species in our analysis, the first subset included data from all species, however we randomly selected 9 individuals and 10 calls per individual, based on the smallest common denominator across datasets (dataset 1, Figure 1). The second was used for analyses investigating differences in results given different sample sizes and number of individuals and consisted of 4 mammalian species representing diverse taxa: Egyptian fruit bat, common marmoset (phees), meerkat, and domestic sheep (dataset 2, Figure 1).

**Figure 1.**
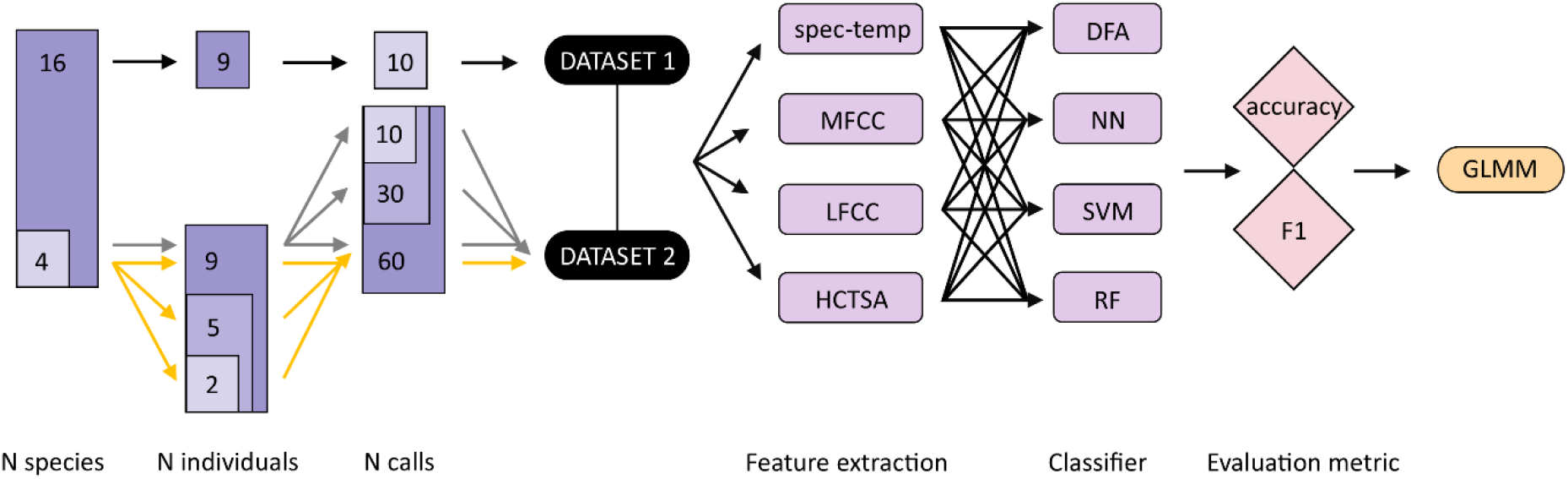
Sequential steps of the methodology used for evaluating the robustness of individual identity estimation in mammal vocalisations. Notations: spec-temp – spectro-temporal features, MFCC – Mel-frequency Cepstral Coefficients, LFCC – Linear frequency cepstral coefficients, HCTSA - Highly Comparative Time Series Analysis, DFA – Discriminant Function Analysis, NN – Neural Networks, SVM – Support Vector Machines, RF – Random Forest, GLMM – Generalised Linear Mixed Effects Model.

#### Feature extraction

Spectrogram features were extracted for all datasets using Raven Pro 1.6.4 (K. Lisa Yang Center for Conservation Bioacoustics, 2024). To limit bias resulting from different researchers processing data, we extracted robust measurements from the calls (Centre Frequency, Frequency 5%, Frequency 25%, Frequency 75%, Frequency 95%, Duration 90%, Duration 50%, Bandwidth 50%, Bandwidth 90% (Charif et al., 2010) which are based on the energy stored within the sound rather than boundaries of a selection (and are therefore less sensitive to user variability), as well as average and aggregate entropy measures (spectro-temporal dataset). Data providers were instructed to provide separate wav files for each call, that were tightly cropped around the call. The whole file was fed into Raven from which measurements were extracted. We also extracted MFCC for all datasets (package BehaviouR (Clink, 2023a)) from the recorded vocalisations (frequency minima and maxima defined separately for each species; MFCC dataset; Supplementary Materials (SM) Table S1). As MFCC have been developed for human speech and some concerns exist about their applicability to non-human animals, especially those vocalising at high frequencies, we also included linear frequency cepstral coefficients (LFCC) in our analysis. These features were extracted using spafe.features.lfcc.lfcc (ver 0.3.3) in Python (ver. 3.13.1), using the same parameters and processing steps as the MFCC function, ensuring directl comparability. In addition, we extracted features using a new approach – the Highly Comparative Time Series Analysis (HCTSA) using the MatLab HCTSA toolbox (Fulcher et al., 2013; Fulcher & Jones, 2017). HCTSA interprets the acoustic waveform as a time series of pressure points, applies a range of time-series analyses to the acoustic data, and generates a matrix of feature measurements. This method has been successfully employed in previous studies for analysing animal vocalisations (Paul et al., 2021; Phaniraj et al., 2023). Consequently, four feature extraction methods (spectro-temporal, MFCC, LFCC and HCTSA; Figure 2) were used and included in further analysis.

**Figure 2.**
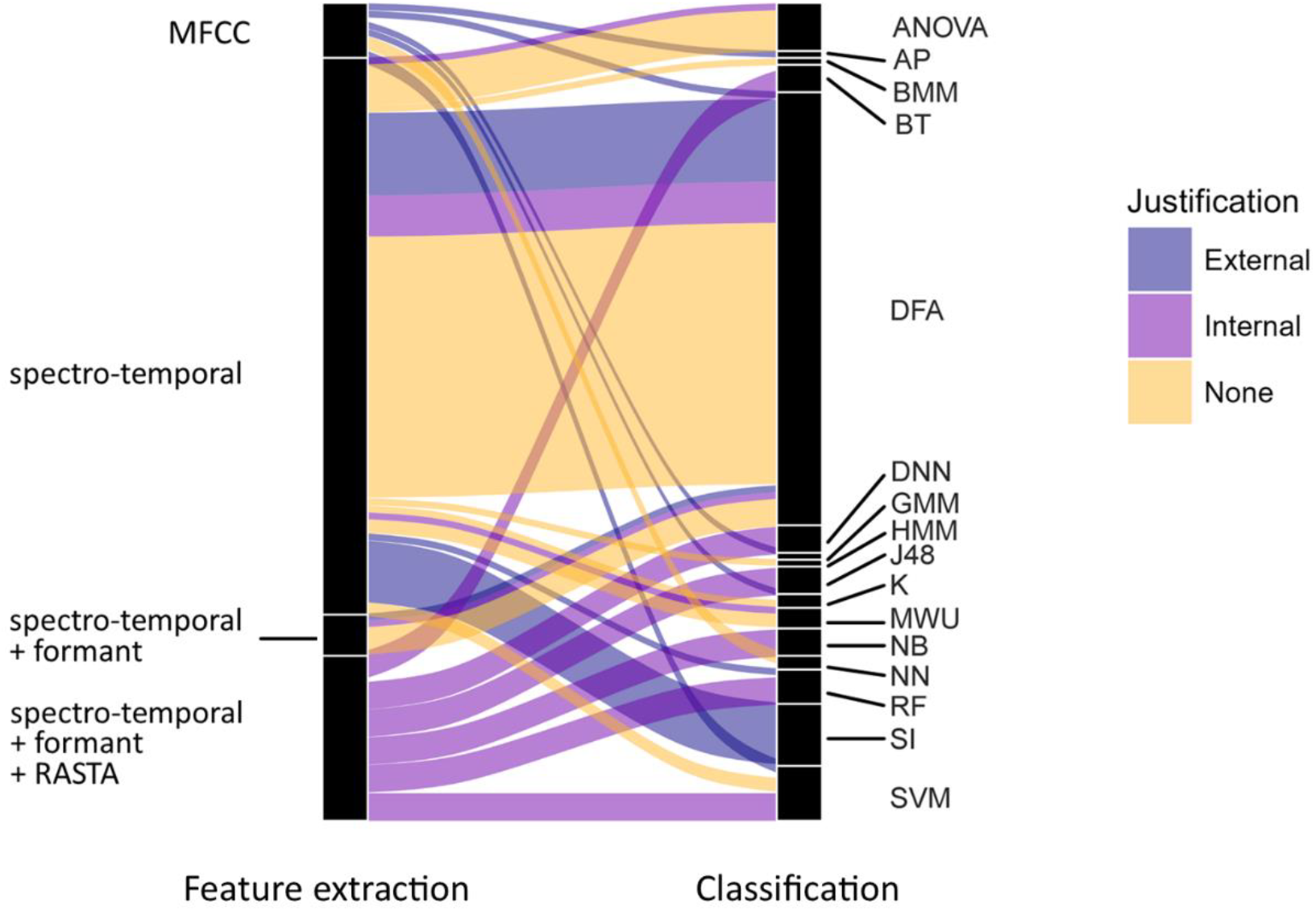
Acoustic feature extraction methods and classifiers used to evaluate individual distinctiveness in mammalian vocalisations, based on a literature review of empirical articles published in years 2000-2022. N articles = 46, N datapoints = 119 (as certain studies encompassed multiple analyses, call types, or species). See SM section 2 for details. Notations: ANOVA – analysis of variance – traditional or multivariate; AP – affinity propagation; BMM – Bayesian functional mixed model; BT – bagged trees; DFA – discriminant function analysis (normal and permutational); DNN – deep neural network; DTW – dynamic time warping; GMM – Gaussian mixture model; HMM – hidden Markov model; K – kmeans or kmedioids; MWU – Mann-Whitney-U test; NB – naïve Bayes; NN - neural network; RF – random forest; SI – stereotypy Index; SVM – support vector machine; MFCC – Mel frequency cepstral coefficients; spectro-temporal – frequency and temporal parameters extracted from the spectrogram, RASTA – relative spectral processing, External justification – based on published work; Internal justification – based on prior experience with the study system.

#### Classification

We applied four commonly used classifiers (discriminant function analysis (DFA), neural networks (NN), support vector machines (SVM), random forests (RF)) to each dataset (spectro-temporal, MFCC, LFCC and HCTSA) for each species, to determine how accurately each one assigns calls to individuals (Figure 2). As the goal was not to obtain the most accurate classifier possible, but to determine how comparably they performed, we did not optimise the classifier to each dataset separately. For all analyses, data were split 70:30 into training and testing sets (within individual and with no overlap), with results reported for testing sets only.

For SVMs, we implemented a radial basis kernel type, using a modified version of the trainSVM function (GibbonR package; Clink, 2023b); the modification split the data without overlap). This function automatically tunes gamma and cost parameters. We ran a RF classifier (randomForest::tuneRF; Liaw & Wiener, 2002) using the optimal mtry found (doBest=TRUE), with 1000 trees used at the tuning step. Neural networks can be affected by collinearity, therefore, we performed a PCA (stats::prcomp; with variables zero-centered and scaled to have unit variance and kept only PCs with eigenvalues>1) prior to running a Model Averaged Neural Network (modified version of the trainNN function; modification as above; GibbonR package (Clink, 2023b)). Before classifying the data using DFA (MASS::lda; Venables & Ripley, 2002), we also conducted a PCA (as above) and the data were normalised (bestNormalize::bestNormalize; Peterson & Cavanaugh, 2020). Our groups were independent and mutually exclusive, and sample size was balanced. Potential collinearity issues were resolved by applying DFA only to data following a PCA. Multivariate normality (MVN::mvn; Korkmaz et al., 2014) and homogeneity of covariances (biotools::boxM; da Silva et al., 2017) were tested. While we tested for the assumptions of a DFA, in order to gauge the robustness of the existing literature’s findings, we extended our analysis to include datasets that did not strictly satisfy the DFA assumptions. To be able to directly compare classifiers we additionally ran the RF and SVM on the PCA and normalised dataset (exactly the same as required and used by the DFA and NN).

Based on the confusion matrices obtained from running the classifiers, we calculated two widely used evaluation metrics: accuracy and F1 scores. Accuracy gives a measure of the proportion of correctly classified samples returned by a classifier, but in some situations can be misleading with regard to the amount of incorrectly classified samples, particularly with imbalanced classes. F1 scores are the harmonic mean of precision and recalland are often more useful in “real world” situations, where classes are imbalanced and there is a high cost associated with incorrect classifications. Here, we have a set of multi-class classification problems with balanced classes (i.e., the same number of calls per individual), so we use accuracy computed as the sum of the diagonal elements of the normalised confusion matrix as this accounts for total number of possible classes (i.e., individuals). We used caret::confusionMatrix (Kuhn & Max, 2008) to compute both accuracy and F1 scores. To account for variation in the results due to initial conditions of the classifiers, we ran 20 iterations (20 seeds) of training for each classifier, each with initial conditions set under a different random seed value.

#### Statistical analyses

All analyses were conducted in R version 4.2.2 (R Core Team, 2022), with linear models fitted using glmmTMB:glmmTMB (Brooks et al., 2017). All P-values we reported are based on likelihood ratio tests (LRT) (Dobson, 2002). For each analysis, we fitted a generalized linear mixed model (GLMM) with beta error distribution and logit link function. Prior to fitting each model, we transformed the response to avoid values being exactly zero or one (Smithson & Verkuilen, 2006). Results presented in the manuscript are model predicted values, that have been back-transformed to the response scale for easier interpretation.

In order to avoid overconfident GLMM estimates and to keep type I error rate at the nominal level of 0.05, we used a maximal random effects structure, as much as possible (Barr et al., 2013; Schielzeth & Forstmeier, 2009) That is, we included random slopes of each the fixed effects and their interactions, where appropriate within each of the random intercepts in each model. For all models, we originally included parameters for the correlations among random slopes and intercepts, but as the models did not converge, we had to exclude them. To include the fixed effect terms as random slopes, we therefore manually dummy coded and centred them to a mean of zero. See below for specific details of each model.

We determined model stability of each model by dropping individual levels of each random factor from the data, one at a time, and comparing the estimates derived from models fitted on the respective subsets with those obtained for the full dataset and found the models to be of acceptable stability (see results).

#### Consistency of classification performance

Our first aim was to estimate how classification accuracy and F1 scores vary depending on the choice of extraction method and classifier. Using a 16 mammalian datasets, with 9 individuals per species and 10 calls per individual, we fitted one GLMM, with accuracy as the response variable and another with F1 as the response variable. In both models, we included classifier (a factor with four levels), extraction method (a factor with four levels) and their interaction as fixed effects and a crossed random effect structure with one random intercept of species (a factor with 16 levels) and another of seed (factor with 20 levels).

As an overall test of the effect of classifier and its interaction with extraction method, and to avoid ‘cryptic multiple testing’ (Forstmeier & Schielzeth, 2011), we compared each model with an appropriate null model lacking the classifier term in the fixed effects but being otherwise identical. We obtained confidence intervals (CIs) of model estimates and fitted values by means of a parametric bootstrap (N=1,000 bootstraps; function simulate of the package glmmTMB; note that 726 and 745 of the bootstrapped models successfully converged for the accuracy and F1 models, respectively). The models were not overdispersed (model dispersion parameters: accuracy model = 1.03; F1 model = 0.93). The samples analysed with these models each comprised a total of 5120 scores for each metric (i.e., 16 mammalian datasets, x 4 extraction methods x 4 classifiers x 20 iterations).

In the analyses described above, we applied a PCA to the data outputted by each of the feature extraction methods before using the DFA and NN classifiers as this is a required step for these classifiers. However, as the RF and SVM classifiers are typically used on the raw data outputted by the feature extraction methods, we did not apply a PCA to the data before using these classifiers. Therefore, to determine whether the RF and SVM classifiers performed similarly with raw data and PCA data, we ran a second set of GLMMs, same as the ones described above, but all classifiers were fed in reduced (PCA) datasets. We compared each model with an appropriate null model lacking the classifier term in the fixed effects but being otherwise identical, and we calculated bootstraps using the same method as above (753 and 751bootstrapped models converged for the accuracy and the F1 models, respectively). The models were not overdispersed (model dispersion parameters: accuracy model = 1.08; F1 model = 1.03). The samples analysed with these models each comprised a total of 5120 scores for each metric (i.e., 16 mammalian datasets, x 4 extraction methods x 4 classifiers x 20 iterations).

#### Effect of sample size of vocalisations: n60 vs n30 vs n10

As high-quality data collected from animals can often be limited in sample size, we also examined how the observed accuracy of the classifiers varied with the sample size (i.e., number of calls per individual) included. We extracted data for which we had 60 calls per individual, for 9 individuals per species. This resulted in a dataset consisting of four species: Egyptian fruit bat, common marmoset (phee), meerkat and domestic sheep. We then created subsets of the data (samples selected randomly, where a different random subset was selected for each seed value) that varied in sample size, with 60 (n60 dataset), 30 (n30 dataset) and 10 (n10 dataset) calls per individual (the number of individuals remained the same at n=9 per species). In the model, we included classifier (a factor with four levels), extraction method (a factor with four levels), sample size (a factor with three levels) and the 3-way interaction between them as a fixed effect. We used a crossed random effect structure, with one random intercept effect of species (a factor with four levels) and another of seed (a factor with 20 levels). Note that the GLMM used in this analysis did not converge with the interactions between the random slope terms, so we removed these interaction terms from the model, but we retained the additive effects of the random slope terms.

As an overall test of the effect of sample size and its interaction with extraction method and classifier, we compared the model with an appropriate null model lacking the sample size term in the fixed effects but being otherwise identical.

We obtained CIs of model estimates and fitted values by means of a parametric bootstrap using glmmTMB::simulate (N=1000 bootstraps; note that only 827 of the bootstrapped models converged). The model was not overdispersed (dispersion parameter=1. 02). The samples analysed with this model comprised a total of 3840 accuracy scores (i.e., 4 species x 4 extraction methods x 4 classifiers x 3 sample size sets x 20 iterations).

#### Effect of number of individuals – 9 vs 5 vs 2 individuals

To determine the consistency of classifier performance when there are fewer possible classes to assign, we ran a similar analysis to the one described above, using subsets of the data restricted in this case by the number of individuals rather than the number of calls per individual. In the first subset, we randomly selected 5 individuals from each of the 4 species, and in the second, we randomly selected 2 individuals. This resulted in 3 datasets per seed, with 9, 5 and 2 individuals per species respectively. Each dataset contained 60 calls per individual, and in each of the 20 iterations, a different set of individuals was randomly selected. Note that the GLMM used in this analysis did not converge with the interactions between the random slope terms, so we removed these interaction terms from the model, but we retained the additive effects of the random slope terms.

As an overall test of the effect of number of individuals and its interaction with extraction method and classifier, we compared the model with an appropriate null model lacking the number of individuals term in the fixed effects but being otherwise identical.

CIs of model estimates were calculated as above (921 of the bootstrapped models for the successfully converged). The model was not overdispersed (dispersion parameters =1.04). The samples analysed with this model comprised a total of 3840 accuracy scores (i.e., 4 species x 4 extraction methods x 4 classifiers x 3 sample size training sets x 20 iterations).

## RESULTS

### LITERATURE REVIEW

Out of the 1000 articles retrieved in our search, we identified 46 distinct papers that were relevant to our study – focusing on empirical investigations of mammalian species incorporating acoustic analysis and evaluating the individual distinctiveness of vocalisations. We obtained a total of 119 observation points, as certain studies encompassed multiple analyses, call types, or species.

We found a considerable diversity in methods used across studies (Figure 2). While a majority of studies relied on spectral features to ascertain the presence of individual distinctiveness (Figure 2), the specific spectral measurements employed varied greatly with 41 unique parameter sets used across studies (Supplementary Materials, dataset 1). Some studies adopted measures targeting the filter-shaped components of the sound, extracting features such as MFCC or formants. Notably, formants were consistently extracted in conjunction with spectral features and were never utilised in isolation. Moreover, a limited number of studies incorporated both source-and filter-related elements of the sound. We found that 24.37% of studies provided external (i.e., citation-based) justification for their approach towards methods used, 27.73% described it as being based on their prior experience with the system (internal justification), and 47.9% provided no justification at all (Figure 2).

Classifiers used to assign a call to an individual varied greatly between studies, encompassing a total of 18 different classifiers (Figure 2). Among these, the most frequently employed were Discriminant Function Analysis (DFA; 52.94%), Analysis of Variance (ANOVA 5.88%), Support Vector Machine (SVM; 6.7%), Random Forest (RF; 4.2%). DFA emerged as the predominant classification method, although it is noteworthy that a majority of authors did not verify, or at least did not report, whether the assumptions of DFA were met – within our literature search, a mere 11.1% of DFA analyses explicitly reported checking the assumptions.

The studies incorporated into our literature analysis encompassed information from nine different mammalian orders, with Primates (43.5%), Carnivora (23.9%), and Chiroptera (6.5%) being the most frequently represented. Furthermore, data were available for 37 genera of mammals (SM Table S2).

### COMPARISON OF FEATURE EXTRACTION AND CLASSIFICATION METHODS

#### Consistency of classification performance

Using acoustic data spanning 16 mammalian datasets we modelled the accuracy and F1 scores obtained from classifiers using different feature extraction methods (Highly Comparative Time Series Analysis (HCTSA), Mel-frequency cepstral coefficients (MFCC), linear frequency cepstral coefficients (LFCC) and spectro-temporal features) as well as classifiers (discriminant function analysis (DFA), neural networks (NN),, random forests (RF), support vector machines (SVM)).

Overall, the full-null model comparisons revealed a clear effect of classifier in each model (accuracy: χ2=334.24, df=12, P<0.001; F1: χ2=396.72, df=12, P<0.001; SM Table S3-S4) and the likelihood ratio tests (LRT) showed that the effects of the interactions between extraction method and classifier were significant (accuracy: χ2=216.92, df=9, P<0.001; F1: χ2=238.95, df=9, P<0.001). For a breakdown of these results by species, see Supplementary Materials (Figure S1, Table S5).

For both evaluation metrics, model results revealed the same general pattern of performance for each combination of extraction method and classifier (Figure 3). Classifiers trained on data extracted using the MFCC feature extraction method all typically provided the highest and most consistent scores (difference between maximum and minimum predicted accuracy values Δ_Accuracy_ = 0.102, Δ_F1_ = 0.114). LFCCs provided comparable results (Δ_Accuracy_ = 0.115, Δ_F1_ = 0.132). A greater variation in the performance of classifiers was observed when using the spectro-temporal dataset (Δ_Accuracy_ = 0.180, Δ_F1_ = 0.227), and to an even greater extent, HCTSA feature extraction methods (Δ_Accuracy_ = 0.582, Δ_F1_ = 0.613). Out of classifiers, random forests resulted in the least amount of variation (Δ_Accuracy_ = 0.194, Δ_F1_ = 0.205) across various extraction methods, followed by SVM: Δ_Accuracy_ = 0.253, Δ_F1_ = 0.276). DFA and NN provided most inconsistent results, with NN having the highest variation (Δ_Accuracy_ = 0.515, Δ_F1_ = 0.542; DFA Δ_Accuracy_ = 0.331, Δ_F1_ = 0.338; Figure 3; SM Table S3, Figure S2).

**Figure 3.**
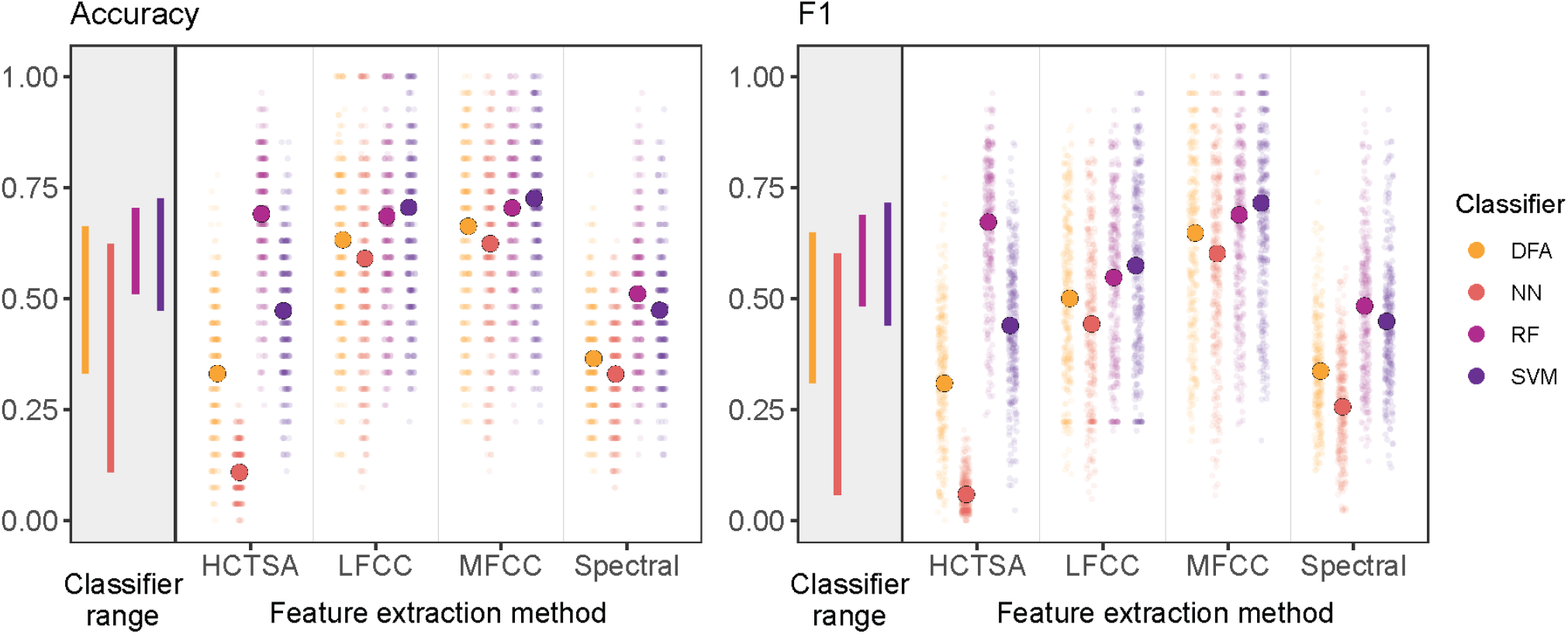
Accuracy and F1 scores of random forest algorithms classifying vocalisations to individual, across 16 mammalian datasets. Observed (smaller points) and GLMM predicted estimates (larger points) for each classifier as plotted across various extraction methods. Vertical lines represent the range of values for each classifier. Dataset consists of vocalisations from 16 species, 9 individuals per species and 10 calls per individual. Values were back transformed to the response scale.

Results conducted on the reduced (PCA) dataset showed similar trends, with MFCC and RF providing more consistent accuracies than the other feature extraction and classification methods (SM Figure S3, Table S6).

#### Effect of sample size of vocalisations on method performance – n60 vs n30 vs n10

Given the common limitations in obtaining high-quality acoustic data from animals, we explored how the sample size affects the accuracy of our classifiers under different feature extraction methods. Specifically, we analysed the performance variation across datasets with different numbers of vocalisations per individual, namely 60 (n60), 30 (n30), and 10 (n10) calls. For this analysis, we used datasets comprising 60 calls from each of the 9 individuals, encompassing 4 distinct species.

The full-null model comparison revealed a clear effect of sample size (number of vocalisations per individual) on accuracy (χ2=1209.247, df=32, P<0.001) and the 3-way interaction between sample size, extraction method and classifier was significant (LRT: χ2=727.663, df=18, P<0.001) (Figure 4A; Accuracy: SM Table S7; F1: SM Figure S4A, Table S8), with the same trends visible as described in the previous section (MFCC and RF providing most consistent results across sample sizes).

**Figure 4.**
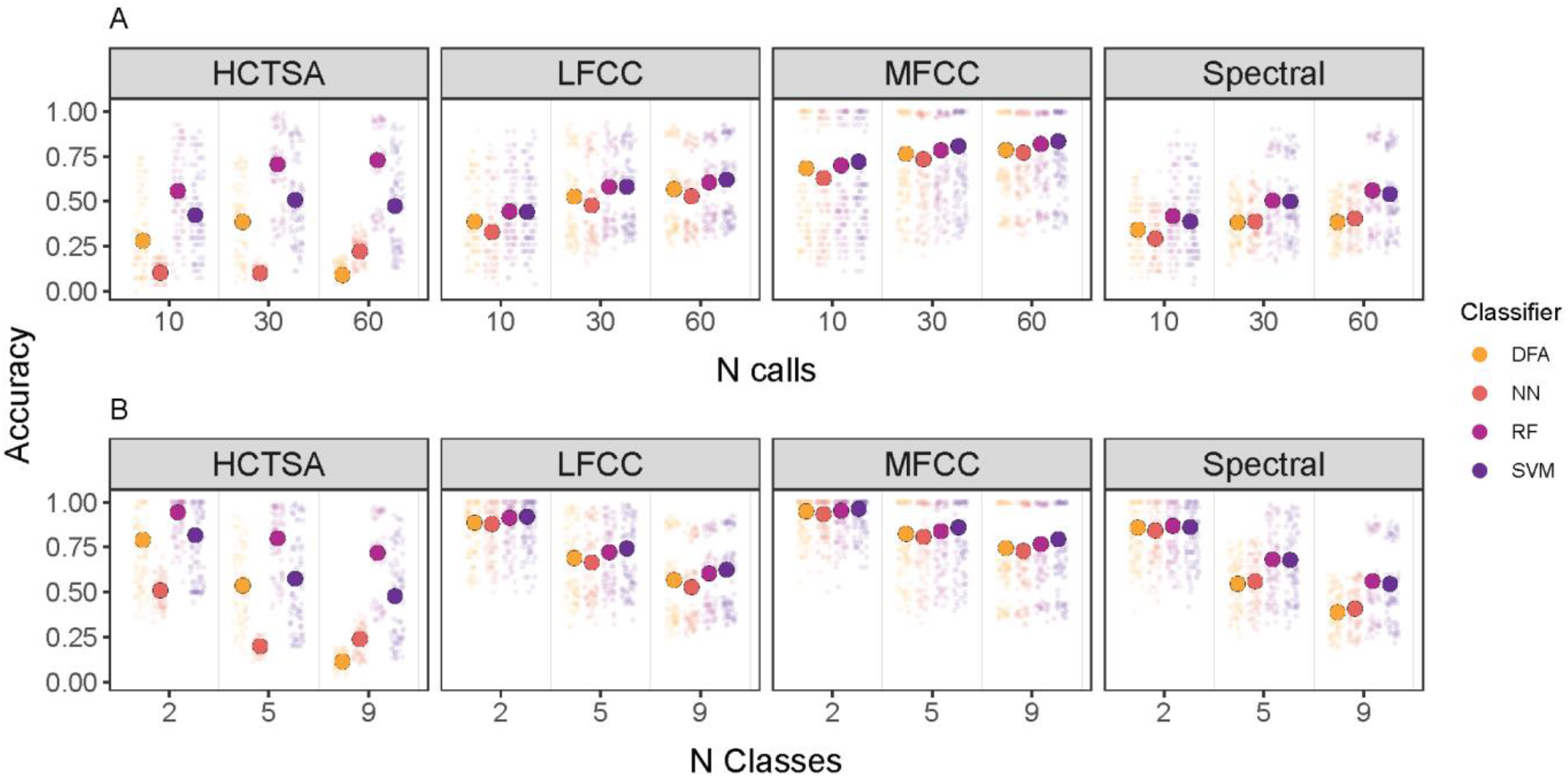
Accuracy scores of random forest algorithms classifying vocalisations to individual for datasets with different A) sample sizes (sample size of 10, 30, or 60 calls per individual, respectively); B) number of classes (2, 5 and 9 individuals per species). Observed (smaller points) and model predicted estimates (larger points) for each classifier as plotted across various extraction methods. Dataset consists of vocalisations from 4 mammalian species. Values were back transformed to the response scale.

For most classifier-extraction method combinations, accuracy increased from n10 to n30, with not much of an increase from n30 to n60 (Figure 4A). The exception to this trend was the HCTSA extraction method paired with the DFA, where increasing the sample size of the training data set to n60 lead to a decrease in accuracy. MFCC datasets typically led to more consistent and better accuracy values than the other extraction methods for all sample sizes, regardless of which classifier they were paired with. Across classification methods, RF performed most consistently across extraction methods in datasets of all sample sizes.

#### Effect of number of individuals – 9 vs 5 vs 2 classes

To evaluate performance with a reduced number of classes, we conducted a similar analysis to the previous one, this time limiting the number of individuals instead of the number of calls per individual. We included three datasets with 9, 5, and 2 individuals per species, each containing 60 calls per individual.

The full-null model comparison revealed a clear effect of the number of individuals on accuracy (χ2=584.791, df=32, P<0.001) and the LRT showed that the 3-way interaction between number of classes, extraction method and classifier was significant (χ2=394.618, df=18, P<0.001). While the overall trends were similar to those reported above (MFCC and RF performed most consistently), we observed more variation in the absolute values of the accuracy achieved by the classifiers within and between sample size groups. Accuracy values were highest and more consistent when a smaller number of individuals was present (Figure 4B; SM, Table S9, Table S10, Figure S4B).

## DISCUSSION

Our literature review confirmed our impression that extraction and classification methods vary greatly across studies investigating the encoding of individual identity in mammalian calls. Authors frequently neglect to substantiate the selection of acoustic features for analysis, utilising a diverse array of classifiers. This results in many findings being isolated, with conclusions that are exclusively species-specific, thus limiting their applicability to a restricted number of researchers. Consequently, the generalizability of results is limited, hindering the derivation of broader evolutionary insights. Here, we assessed the comparability of various combinations of feature extraction and classification methods commonly used for the identification of individual vocal signature, aiming to establish a guideline for future research endeavours that facilitate standardised inter-taxa comparisons and contribute to larger-scale analyses and overarching conclusions on vocal individuality. We investigated the accuracy of these methods in assigning identity across 16 call types from 14 different mammalian species. Our findings showed that using Mel-frequency cepstral coefficients to extract acoustic features, or random forests as classifiers provides relatively consistent results. We therefore recommend using these methods in future investigations to enhance comparability of results across species.Obtained accuracy values varied significantly depending on the employed feature extraction methods and classifiers. Results obtained from datasets using MFCC (describing the spectral envelope of the sound) as the extraction method were most robust and provided the most consistent results across different classifiers used. As filter-related acoustic parameters (such as MFCC) of the vocalisations reflect individual morphology and cannot be easily modified without a flexible vocal tract, they tend to be a more stable and reliable cue and are therefore more predictable (Taylor et al., 2016). This information can be used by animals directly for individual recognition, or to make the recognition process more efficient by distinguishing broader categories of individuals first (e.g., size, sex, age) and then identifying the individual. Parameters related to the spectral envelope have been extensively used for voice recognition in humans and have been shown to be an important cue for recognition in other non-human primates (Clink et al., 2019, 2020; Gamba et al., 2012; Mielke & Zuberbühler, 2013) as well as other taxa (Townsend et al., 2014). The results obtained using LFCC values were highly comparable to those derived from MFCC across all species, highlighting the suitability of these methods for analysing vocalizations of non-human animals, including those that produce higher-frequency calls. While our study did not specifically focus on extreme frequency ranges, and only a small portion of our data included ultrasounds, species-specific plots (Supplementary Materials, Figure S1) indicate that MFCC and LFCC performed similarly well.

Results obtained when using HCTSA datasets were the most variable. This can be attributed, at least in part, to the very large number of features returned by this extraction method. A challenge arises when limited sample sizes intersect with a large number of features, potentially impeding the algorithm’s accuracy during classification (usually due to overfitting). Conversely, a lower number of features may offer less information per datapoint during classification, which may be the reason for the lower accuracy values from the spectro-temporal dataset. We deliberately refrained from constraining the number of features across datasets, recognising it as an intrinsic characteristic of the extraction method that warrants consideration in the choice of methodology used for each study. Additionally, it is crucial to acknowledge that a larger number of features facilitates a more detailed description of the sound, although it is difficult to say if all of these features are biologically relevant. The number of features to be used for classification thus requires careful consideration on a case-by-case basis, contingent upon the specific objectives and constraints of each study.

Among the various classifiers, RF consistently yielded the least variable results across diverse datasets (SVM produced relatively high accuracy values, yet were more slightly variable then RF results across different extraction methods). This can be attributed to the algorithm’s ease of use, requiring few assumptions of the data. Conversely, DFA (commonly employed in previous studies, 52.94% of the cases in our literature review) and NN exhibited the lowest accuracy values. Results derived from DFA are only reliable when stringent assumptions are met. Given the limited availability of studies providing information on assumption testing, there is a high likelihood that results derived from DFA may lack reliability, precluding meaningful comparisons with other studies and warranting caution in future analyses.

The lower accuracy values obtained from the DFA and NN classifiers could be partially due to the reduced number of variables (PCA-transformed dataset), limiting the information available to the algorithm. However, even when all classifiers were evaluated using a PCA-transformed dataset, RF still produced the most consistent results. Regardless, the need for dimensionality reduction in NN and DFA (but not in RF and SVM) is an inherent characteristic of these algorithms and cannot be disentangled from their performance. Our objective was not to determine the best classifier in general, but rather the most consistent one for this specific task. If a classifier performs best because it does not require dimensionality reduction, that advantage remains valid for this application. Moreover, avoiding dimensionality reduction may simplify the analysis and reduce potential sources of error. Thus, whether due to its ability to incorporate all raw variables or other factors, RF consistently delivers the most consistent and reliable results and could be effectively used for species comparisons.

In our exploration of machine learning approaches for individual identification in animal vocalisations, we encountered considerable variation in results contingent upon the feature extraction and classification methods used. This variability prompts a critical examination of how results from different studies are compared. Each study’s unique way of processing and analysing vocal data means that outcomes may not be directly comparable. Such discrepancies highlight the importance of contextualising findings within their methodological frameworks. While this does not invalidate the results, it does caution against overly simplistic cross-study comparisons. Researchers must recognise that differences in feature extraction and processing approaches can significantly influence the outcomes, thus making it essential to approach inter-study comparisons with a nuanced understanding of the underlying methodologies.

We investigated the effect of different sample sizes on the obtained accuracies. Overall trends of accuracy values remained consistent, even when reducing sample sizes to 10 calls per individual, albeit with slightly lower values compared to n30 and n60 datasets. Given the inherent challenges associated with acquiring acoustic data, particularly from wild animals, the optimal approach, if feasible, appears to be targeting approximately 30 calls per individual. However, it seems that there is no compelling need to further increase the sample size beyond this threshold. Importantly, a sample size of 10 calls per individual presents trends comparable to those obtained with larger sample sizes.

We varied the number of individuals, to be classified, and found that the main trends were similar to those described above, with MFCCs and RF having least variation in accuracy values. As expected, since the task of classifying vocalisations to fewer groups is easier, we observed greater and more consistent values for the smaller number of individuals, with the accuracy values for two individuals being notable greater than for five or nine individuals (with little difference between the latter two). This synthesis provides a comprehensive overview of the current landscape of machine learning applications in individual identity classification of animal vocalisations. While recognising the importance of tailoring analyses to species-specific characteristics and the biological questions posed, our findings indicate that employing Mel-frequency cepstral coefficients and random forests enhances comparability across species. As an initial step in establishing guidelines for processing mammalian acoustic data, where possible, we recommend including these in analyses, to facilitate future comparative studies. We encourage authors to consider these analyses in conjunction with species- and question-specific approaches, contributing to the creation of a shared foundation (however, it is important to acknowledge that these results capture only machine-detectable differences between vocalisations, and in many cases, the biological relevance of these differences for the studied species remains unknown). To support this approach, we provided the code for these analyses (SM, section 1). We hope that our contribution will facilitate a shift towards methodological consistency in the field of bioacoustics, arming future researchers with the knowledge to perform analyses that will produce the directly comparable outputs (both across species, and across studies within species) necessary to answer the most challenging research questions.

## ACKNOWLEDGEMENTS

We thank Dr Roger Mundry for sharing R functions providing various diagnostics for evaluating model assumptions and stability and setting up an appropriate random slopes structure for (G)LMMs.

## Funding

KW, SKW, NF, MBM, NP and JMB were partially or entirely funded by the NCCR “Evolving Language”, Swiss National Science Foundation Agreement #51NF40_180888. CF thanks the SFB1528 “Cognition of interactions” for discussions.

## AUTHOR CONTRIBUTIONS

Kaja Wierucka conceived the ideas and designed methodology; Claudia Fichtel, Julian León, Stephan Leu, Elodie Briefer, Marta Manser, and Marina Scheumann contributed the data; Kaja Wierucka prepared and consolidated the data; Kaja Wierucka and Derek Murphy analysed the data; Nikhil Phaniraj and Nikola Falk scored the literature review; Kaja Wierucka led the writing of the manuscript. All authors contributed critically to the drafts and gave final approval for publication.

## Notes

### Competing Interest Statement

The authors have declared no competing interest.

### Summary of Updates

Minor changes to figures, LFCC extraction method added

