## Supplementary material for "Same data, different results? Machine learning approaches in bioacoustics": SM

### 1. DATA SOURCES

Data used in this study have been collected for other studies, with data contributors serving as co-authors.

Data collection methods or sources are detailed below.

Alpaca (*Vicugna pacos*): Data taken from Wierucka & Leu (2024).

Domestic cat (*Felis catus*): Data originated from the sound archive of the Institute of Zoology, University of Veterinary Medicine Hannover, originally published in Scheumann et al. (2012)

Egyptian fruit bat (*Rousettus aegyptiacus*): Data taken from Prat et al. (2017).

Domestic goat (*Capra aegagrus hircus*): Data taken from Briefer et al. (2015).

Domestic horse (*Equus caballus*): Data taken from Briefer et al. (2015).

Sooty mangabey (*Cercocebus atys*): Data taken from León et al. (2022).

Common marmoset (*Callithrix jacchus*): We collected vocalisations from adult common marmosets housed at the University of Zürich from January to August 2022. The animals had access to water freely throughout the day. Their diet consisted of vitamin-enriched mash and/or fresh fruits and vegetables, provided at least twice daily: once in the morning and again at noon. In the afternoon, they were given additional protein-rich food such as insects, gum, boiled eggs, or cheese. The marmosets were housed in separate heated indoor enclosures and had access to outdoor areas, either in groups or pairs. All procedures followed Swiss legislation and were approved by Zürich's cantonal veterinary office (license ZH223/16). Recording sessions involved two marmosets, placed in adjacent wire cages. Each recording session lasted for 15 minutes. For audio recordings, we utilized two CM16/CMPA Avisoft Bioacoustics condenser microphones. Caller identity was annotated in real-time using Avisoft Recorder. Phees and trills were manually extracted using Avisoft-SASLab Pro ver. 5.3.01.

Meerkat (*Suricata suricatta*): Recordings of meerkats were collected within the Kalahari Meerkat Project organized by the Kalahari Research Centre, located at the Northern Cape in South Africa (26°58'S, 21°49'E) as part of long-term data collection(Clutton-Brock & Manser, 2016). All individuals in the population were tagged with subcutaneous transponders and were dye-marked in order to facilitate identification in the field(Jordan et al., 2007). Meerkats were wild, but habituated to human presence, allowing observers to follow meerkats with a directional microphone at close distance (0.5 - 2 meters). The recordings, which were collected from 2014 until 2020, were recorded with a Marantz PMD-670 solid-state recorder (Marantz Japan Inc.; sampling frequency 48 kHz or 44.1 kHz, 16 bits) attached to a directional microphone (Sennheiser ME66/K6, Sennheiser Electronic Corp., Old Lyme, CT, U.S.A.). Original sound files were cut and processed in the course of various projects using Adobe Audition and PRAAT software(Boersma & Weenink, 2023). We extracted at least 110 contact calls per individual of 20 adult meerkats (10 male/10 female) produced during the foraging contexts (Doolan & MacDonald, 1996; Townsend et al., 2011) over various events (3 to 14).

Grey mouse lemur (*Microcebus murinus*): Data originated from the sound archive of the Institute of Zoology, University of Veterinary Medicine Hannover, originally published in Leliveld et al. (2011), Scheumann & Zimmermann (2008).

Domestic pig (*Sus scrofa domesticus*): Data taken from Briefer et al. (2019).

Redfronted lemur (*Eulemur rufifrons*): Data taken from Fichtel & Kappeler (2002), Fichtel & Hammerschmidt (2002).

Domestic sheep (*Ovis aries*): The sheep vocalisations were recorded as part of a previously carried out and unpublished playback experiment. Domestic adult Swiss female sheep, housed at AgroVet-Strickhof (Lindau, Switzerland), were recorded in social isolation as follows; sheep were led alone from their home pen to the experimental arena, situated in an adjacent room, where one bucket of food was present. The subject was then left there for 5min and brought back to her home pen afterwards. This procedure was repeated each day for five consecutive days. All sheep were recorded at distances of 3 to 5 m using a Sennheiser MKH-67 directional microphone (frequency response 40 Hz to 20 000 Hz; max SPL 125 dB at 1 kHz), connected to a Marantz PMD-661 numeric recorder, at a sampling rate of 44.1 kHz. Sounds were then imported into a computer at a sampling rate of 44.1 kHz and saved in WAV format at 16-bit amplitude resolution. All good quality bleats (i.e., low level of background noise), as visualised on a spectrogram in Praat v.5.3.41 (FFT method, window length = 0.03 s, time steps = 1000, frequency steps = 250, Gaussian window shape, dynamic range = 60 dB) were then extracted and saved as individual WAV files and stored in a database. All experiments were carried out in accordance with the current laws of Switzerland. This study was approved by the Cantonal authority (approval number ZH226/17; Switzerland).

Squirrel monkeys (*Saimiri sciureus*): Data taken from Jurgens et al. (1979) and Fichtel et al. (2001).

Tree shrews (*Tupaia belangeri*): Data originated from the sound archive of the Institute of Zoology, University of Veterinary Medicine Hannover, originally published in Schehka & Zimmermann (2009).

### 2. SUPPLEMENTARY RESULTS

All results presented in the main text and figures were back transformed to the response scale. Here we present both model estimates on the logit link scale, and fitted values from the models, backtransformed to the response scale.

**Table S1. Range of frequencies used for MFCC (and LFCC) analysis for each species/call type.** Hann window was used with a 50% overlap. Notations: Fmin – minimum frequency, Fmax – maximum frequency

| Species | Calltype | MFCC Fmin [Hz] | MFCC Fmax [Hz] | Window size [samples] |
|---|---|---|---|---|
| alpaca | hum | 50 | 5000 | 1300 |
| domestic cat | isolation call of kitten | 0 | 22000 | 512 |
| Egyptian fruit bat | social call | 2000 | 70000 | 800 |
| domestic goat | | 0 | 8000 | 1500 |
| domestic horse | whinny | 0 | 22000 | 1000 |
| sooty mangabey | grunt | 0 | 2500 | 700 |
| common marmoset | phee | 4000 | 21000 | 400 |
| common marmoset | trill | 4000 | 25000 | 700 |
| meerkat | close call | 0 | 4000 | 600 |
| grey mouse lemur | tsak | 4000 | 90000 | 300 |
| grey mouse lemur | whistle | 6000 | 80000 | 200 |
| domestic pig | grunt | 0 | 8000 | 900 |
| Red-fronted lemur | woof | 0 | 16000 | 400 |
| domestic sheep | bleats | 0 | 10000 | 3000 |
| squirrel monkey | shrieks | 0 | 8000 | 600 |
| tree shrew | chatter | 0 | 10000 | 600 |

**Table S2. Distribution of taxa occurring in scientific articles investigating individual distinctiveness of animal vocalisations based on acoustic features (literature published in years 2000-2022).** A – distribution of orders, B – distribution of genera.

**A.**

| Order | Frequency | Percent |
|---|---|---|
| *Artiodactyla* | 5 | 10.87 |
| *Carnivora* | 11 | 23.91 |
| *Chiroptera* | 3 | 6.52 |
| *Lagomorpha* | 1 | 2.17 |
| *Perissodactyla* | 1 | 2.17 |
| *Primates* | 20 | 43.48 |
| *Proboscidea* | 3 | 6.52 |
| *Rodentia* | 2 | 4.35 |

**B.**

| Genus | Frequency | Percent |
|---|---|---|
| *Arctocephalus* | 1 | 2.17 |
| *Callithrix* | 2 | 4.35 |
| *Canis* | 2 | 4.35 |
| *Carlito* | 1 | 2.17 |
| *Cebus* | 1 | 2.17 |
| *Cercocebus* | 1 | 2.17 |
| *Cercopithecus* | 2 | 4.35 |
| *Cervus* | 1 | 2.17 |
| *Dama* | 1 | 2.17 |
| *Desmodus* | 1 | 2.17 |
| *Diaemus* | 1 | 2.17 |
| *Elephas* | 1 | 2.17 |
| *Hydrurga* | 6 | 13.04 |
| *Hylobates* | 1 | 2.17 |
| *Leptonychotes* | 1 | 2.17 |
| *Loxodonta* | 1 | 2.17 |
| *Lycaon* | 1 | 2.17 |
| *Macaca* | 1 | 2.17 |
| *Microcebus* | 1 | 2.17 |
| *Neophoca* | 1 | 2.17 |
| *Ochotona* | 1 | 2.17 |
| *Pan* | 1 | 2.17 |
| *Panthera* | 1 | 2.17 |
| *Papio* | 2 | 4.35 |
| *Phoca* | 1 | 2.17 |
| *Physeter* | 2 | 4.35 |
| *Pteronura* | 1 | 2.17 |
| *Pteropus* | 1 | 2.17 |
| *Simias* | 1 | 2.17 |
| *Spermophilus* | 1 | 2.17 |
| *Suricata* | 1 | 2.17 |
| *Tadarida* | 1 | 2.17 |
| *Tamiasciurus* | 1 | 2.17 |
| *Tapirus* | 1 | 2.17 |
| *Tarsius* | 1 | 2.17 |
| *Tursiops* | 1 | 2.17 |

**Table S3. Model results (accuracy) for 16 mammalian data sets.**

Table S3a. Results of the model with accuracy as the response variable for data including 16 mammalian data sets with n=10 calls per individual (estimates, together with standard errors (SE), confidence limits (Cl), as well as minimum and maximum of model estimates obtained when excluding individual levels of species and seed one at a time).

| Term | Estimate | SE | Lower CI | Upper CI | Min | Max |
|---|---|---|---|---|---|---|
| intercept | -0.705 | 0.191 | -1.086 | -0.544 | -0.768 | -0.637 |
| extraction (LFCC) | 1.245 | 0.283 | 0.483 | 1.139 | 1.015 | 1.340 |
| extraction (MFCC) | 1.379 | 0.181 | 1.072 | 1.792 | 1.258 | 1.450 |
| extraction (spectral) | 0.149 | 0.094 | -0.059 | 0.338 | 0.102 | 0.202 |
| classifier (NN) | -1.404 | 0.068 | -2.100 | -1.842 | -1.477 | -1.349 |
| classifier (RF) | 1.506 | 0.058 | 1.410 | 1.645 | 1.456 | 1.558 |
| classifier (SVM) | 0.593 | 0.051 | 0.465 | 0.664 | 0.556 | 0.642 |
| extraction (LFCC):classifier (NN) | 1.228 | 0.122 | 1.541 | 1.947 | 1.150 | 1.302 |
| extraction (MFCC):classifier (NN) | 1.234 | 0.107 | 1.597 | 1.946 | 1.176 | 1.310 |
| extraction (spectral):classifier (NN) | 1.251 | 0.109 | 1.388 | 1.779 | 1.183 | 1.310 |
| extraction (LFCC):classifier (RF) | -1.269 | 0.102 | -1.556 | -1.122 | -1.321 | -1.196 |
| extraction (MFCC):classifier (RF) | -1.313 | 0.109 | -1.557 | -1.112 | -1.368 | -1.252 |
| extraction (spectral):classifier (RF) | -0.908 | 0.084 | -1.093 | -0.744 | -0.967 | -0.840 |
| extraction (LFCC):classifier (SVM) | -0.263 | 0.072 | -0.417 | -0.101 | -0.311 | -0.225 |
| extraction (MFCC):classifier (SVM) | -0.298 | 0.072 | -0.387 | -0.116 | -0.345 | -0.260 |
| extraction (spectral):classifier (SVM) | -0.144 | 0.068 | -0.216 | 0.032 | -0.197 | -0.097 |

Table S3b. Fitted values and upper and lower confidence limits (Cl) on the response scale from the model with overall accuracy as the response for data including 16 different species with n=10 calls per individual.

| Extraction method | Classifier | Estimate | Lower CI | Upper CI |
|---|---|---|---|---|
| HCTSA | DFA | 0.331 | 0.252 | 0.422 |
| HCTSA | NN | 0.108 | 0.067 | 0.169 |
| HCTSA | RF | 0.690 | 0.574 | 0.787 |
| HCTSA | SVM | 0.472 | 0.356 | 0.591 |
| LFCC | DFA | 0.632 | 0.391 | 0.813 |
| LFCC | NN | 0.590 | 0.270 | 0.842 |
| LFCC | RF | 0.685 | 0.369 | 0.883 |
| LFCC | SVM | 0.705 | 0.412 | 0.884 |
| MFCC | DFA | 0.662 | 0.482 | 0.808 |
| MFCC | NN | 0.623 | 0.360 | 0.835 |
| MFCC | RF | 0.704 | 0.448 | 0.876 |
| MFCC | SVM | 0.725 | 0.496 | 0.877 |
| spectral | DFA | 0.364 | 0.247 | 0.505 |
| spectral | NN | 0.33 | 0.162 | 0.553 |
| spectral | RF | 0.51 | 0.308 | 0.709 |
| spectral | SVM | 0.473 | 0.308 | 0.667 |

**Table S4. Model results (F1) for 16 species.**

Table S4a. Results of the model with F1 as the response variable for data including 16 mammalian data sets with n=10 calls per individual (estimates, together with standard errors (SE), confidence limits, as well as minimum and maximum of model estimates obtained when excluding individual levels of species and seed one at a time).

| Term | Estimate | SE | Lower CI | Upper CI | Min | Max |
|---|---|---|---|---|---|---|
| intercept | -0.807 | 0.141 | -1.086 | -0.544 | -0.871 | -0.741 |
| extraction (LFCC) | 0.806 | 0.174 | 0.483 | 1.139 | 0.729 | 0.937 |
| extraction (MFCC) | 1.414 | 0.183 | 1.072 | 1.792 | 1.291 | 1.49 |
| extraction (spectral) | 0.127 | 0.101 | -0.059 | 0.338 | 0.079 | 0.181 |
| classifier (NN) | -1.971 | 0.068 | -2.1 | -1.842 | -2.041 | -1.92 |
| classifier (RF) | 1.524 | 0.061 | 1.41 | 1.645 | 1.474 | 1.578 |
| classifier (SVM) | 0.56 | 0.05 | 0.465 | 0.664 | 0.53 | 0.613 |
| extraction (LFCC):classifier (NN) | 1.74 | 0.11 | 1.541 | 1.947 | 1.673 | 1.817 |
| extraction (MFCC):classifier (NN) | 1.774 | 0.094 | 1.597 | 1.946 | 1.717 | 1.848 |
| extraction (spectral):classifier (NN) | 1.582 | 0.101 | 1.388 | 1.779 | 1.517 | 1.647 |
| extraction (LFCC):classifier (RF) | -1.335 | 0.112 | -1.556 | -1.122 | -1.388 | -1.271 |
| extraction (MFCC):classifier (RF) | -1.34 | 0.114 | -1.557 | -1.112 | -1.391 | -1.271 |
| extraction (spectral):classifier (RF) | -0.912 | 0.088 | -1.093 | -0.744 | -0.972 | -0.843 |
| extraction (LFCC):classifier (SVM) | -0.262 | 0.08 | -0.417 | -0.101 | -0.313 | -0.214 |
| extraction (MFCC):classifier (SVM) | -0.248 | 0.068 | -0.387 | -0.116 | -0.295 | -0.205 |
| extraction (spectral):classifier (SVM) | -0.088 | 0.067 | -0.216 | 0.032 | -0.139 | -0.045 |

Table S4b. Fitted values and upper and lower confidence limits (Cl) on the response scale from the model with F1 as the response for data including 16 different species with n=10 calls per individual.

| Extraction method | Classifier | Estimate | Lower CI | Upper CI |
|---|---|---|---|---|
| HCTSA | DFA | 0.309 | 0.252 | 0.367 |
| HCTSA | NN | 0.059 | 0.04 | 0.084 |
| HCTSA | RF | 0.672 | 0.58 | 0.75 |
| HCTSA | SVM | 0.439 | 0.35 | 0.53 |
| LFCC | DFA | 0.5 | 0.354 | 0.645 |
| LFCC | NN | 0.442 | 0.238 | 0.668 |
| LFCC | RF | 0.547 | 0.321 | 0.754 |
| LFCC | SVM | 0.574 | 0.365 | 0.761 |
| MFCC | DFA | 0.647 | 0.497 | 0.777 |
| MFCC | NN | 0.601 | 0.374 | 0.794 |
| MFCC | RF | 0.688 | 0.46 | 0.856 |
| MFCC | SVM | 0.715 | 0.516 | 0.858 |
| spectral | DFA | 0.336 | 0.241 | 0.449 |
| spectral | NN | 0.256 | 0.135 | 0.433 |
| spectral | RF | 0.483 | 0.304 | 0.667 |
| spectral | SVM | 0.448 | 0.304 | 0.62 |

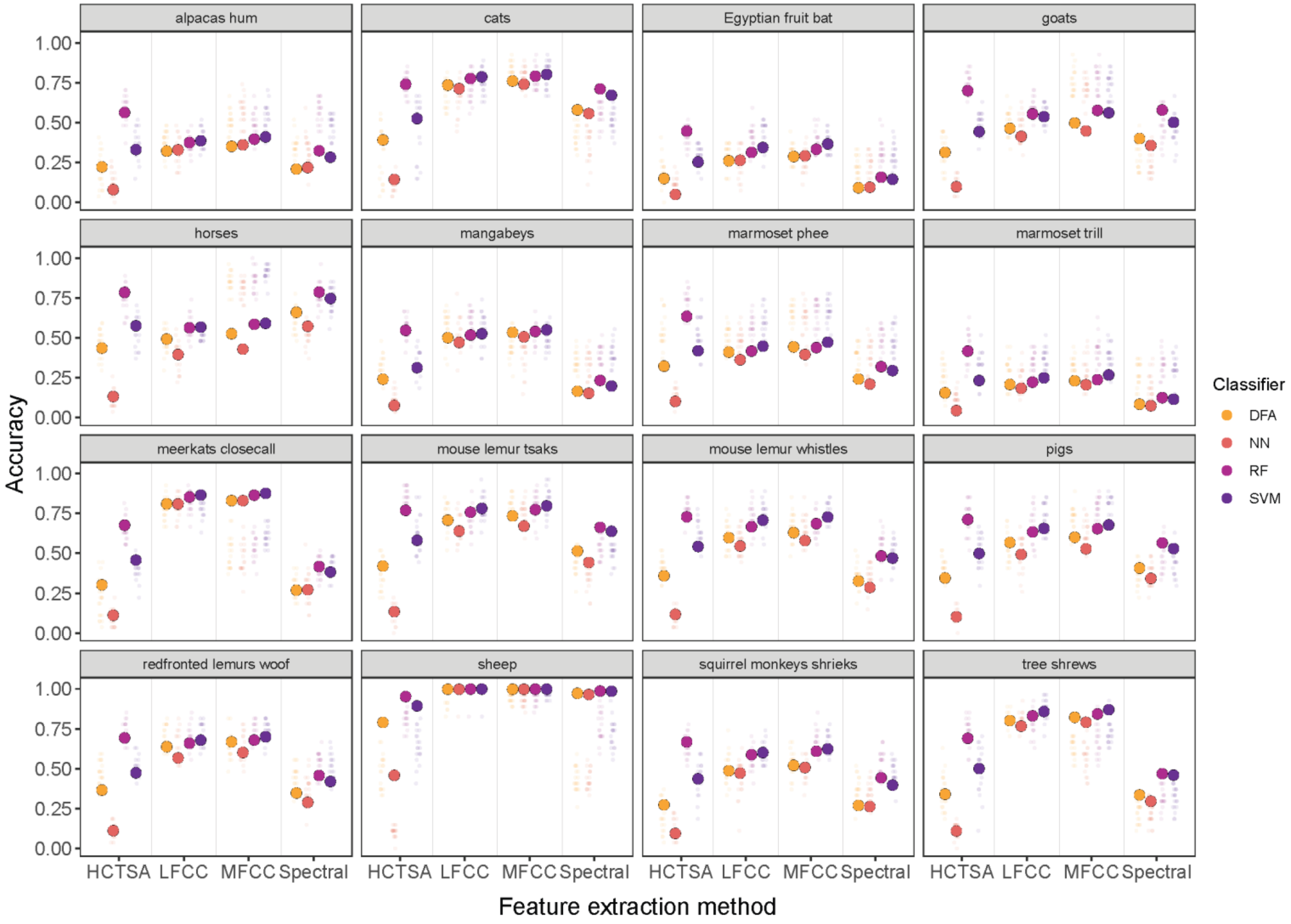


**Figure S1. Observed (smaller points) and model predicted estimates (larger points) of accuracy for each tested species.** Data includes 9 individuals per species and 10 calls per individual. Values were back transformed to the response scale.

**Table S5. Estimated standard deviations (SD) for the contribution of the random effects to the accuracy model (16 mammalian data sets, 9 individuals per species, 10 calls per individual).** The intercept denotes a random intercept effect (Extraction HCTSA and Classification DFA), other entries denote random slope effects.

| Term | effect* | SD |
|---|---|---|
| Species | (Intercept) | 0.674 |
| Species | Extraction LFCC | 1.114 |
| Species | Extraction MFCC | 0.699 |
| Species | Extraction Spectral | 0.328 |
| Species | Classifier NN | 0.135 |
| Species | Classifier RF | 0.125 |
| Species | Classifier SVM | 0.132 |
| Species | Extraction LFCC:Classifier NN | 0.444 |
| Species | Extraction LFCC:Classifier RF | 0.364 |
| Species | Extraction LFCC:Classifier SVM | 0.222 |
| Species | Extraction MFCC:Classifier NN | 0.375 |
| Species | Extraction MFCC:Classifier RF | 0.39 |
| Species | Extraction MFCC:Classifier SVM | 0.222 |
| Species | Extraction Spectral:Classifier NN | 0.381 |
| Species | Extraction Spectral:Classifier RF | 0.287 |
| Species | Extraction Spectral:Classifier SVM | 0.212 |
| Seed | (Intercept) | 0.027 |
| Seed | Extraction LFCC | <0.001 |
| Seed | Extraction MFCC | <0.001 |
| Seed | Extraction Spectral | <0.001 |
| Seed | Classifier NN | 0.056 |
| Seed | Classifier RF | <0.001 |
| Seed | Classifier SVM | 0.013 |
| Seed | Extraction LFCC:Classifier NN | <0.001 |
| Seed | Extraction LFCC:Classifier RF | 0.05 |
| Seed | Extraction LFCC:Classifier SVM | <0.001 |
| Seed | Extraction MFCC:Classifier NN | <0.001 |
| Seed | Extraction MFCC:Classifier RF | 0.049 |
| Seed | Extraction MFCC:Classifier SVM | <0.001 |
| Seed | Extraction Spectral:Classifier NN | 0.093 |
| Seed | Extraction Spectral:Classifier RF | <0.001 |
| Seed | Extraction Spectral:Classifier SVM | <0.001 |

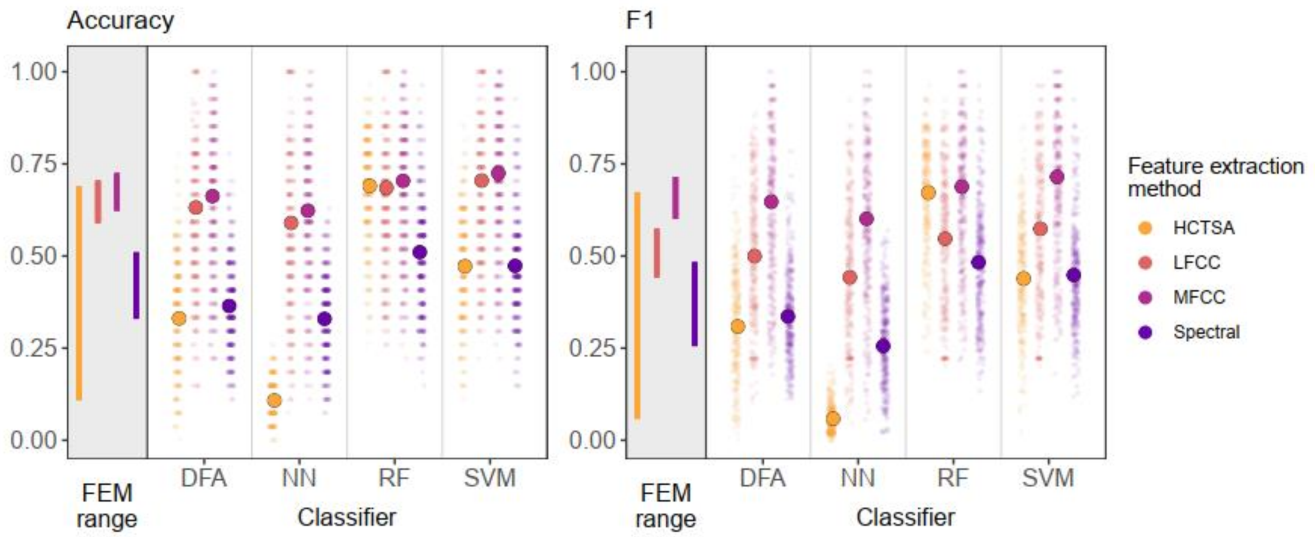


**Figure S2. Accuracy and F1 scores of machine learning algorithms classifying vocalisations to individual, across 16 mammalian species.** Observed (smaller points) and GLMM predicted estimates (larger points) for each extraction method as plotted across various classifiers. Vertical lines represent the range of values for each feature extraction method (FEM). Dataset consists of vocalisations from 16 mammalian data sets, 9 individuals per species and 10 calls per individual. Values were back transformed to the response scale. Note that this is the same figure as in figure 3 of the main text, but with the axes for classifier and feature extraction method flipped to better visualise the trends across classifiers.

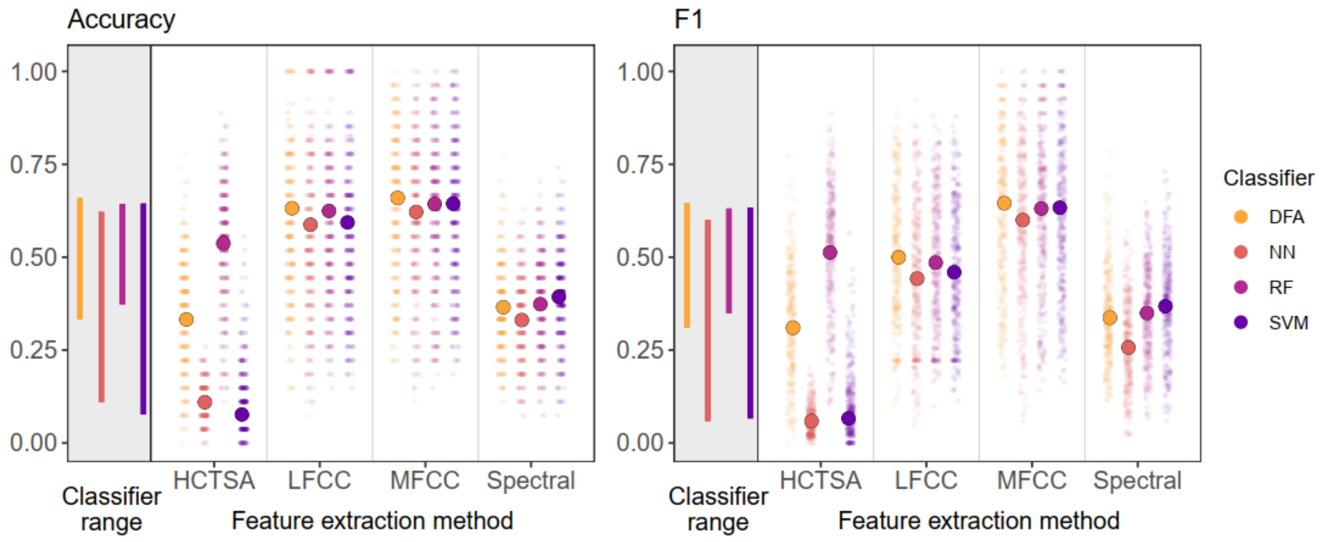


**Figure S3. Accuracy and F1 scores of machine learning algorithms classifying vocalisations to individual, across 16 mammalian species, using only PCA datasets.** Observed (smaller points) and GLMM predicted estimates (larger points) for each classifier as plotted across various extraction methods. Vertical lines represent the range of values for each classifier. Dataset consists of vocalisations from 16 mammalian data sets, 9 individuals per species and 10 calls per individual. Values were back transformed to the response scale.

**Table S6. Model results (accuracy) for 16 species, PCA datasets.**

**Table S6a:** Results of the model with accuracy as the response variable for data including 16 different species with n=10 calls per individual (estimates, together with standard errors (SE), confidence limits (Cl), as well as minimum and maximum of model estimates obtained when excluding individual levels of species and seed one at a time), using only PCA data.

| Term | Estimate | SE | Lower CI | Upper CI | Min | Max |
|---|---|---|---|---|---|---|
| intercept | -0.698 | 0.184 | -1.041 | -0.359 | -0.76 | -0.63 |
| extraction (LFCC) | 1.237 | 0.296 | 0.661 | 1.832 | 1.003 | 1.333 |
| extraction (MFCC) | 1.359 | 0.199 | 0.969 | 1.756 | 1.244 | 1.428 |
| extraction (spectral) | 0.146 | 0.086 | -0.034 | 0.307 | 0.1 | 0.2 |
| classifier (NN) | -1.396 | 0.062 | -1.512 | -1.28 | -1.473 | -1.347 |
| classifier (RF) | 0.847 | 0.053 | 0.742 | 0.946 | 0.802 | 0.891 |
| classifier (SVM) | -1.79 | 0.075 | -1.94 | -1.648 | -1.863 | -1.722 |
| extraction (LFCC):classifier (NN) | 1.21 | 0.104 | 0.998 | 1.419 | 1.151 | 1.286 |
| extraction (MFCC):classifier (NN) | 1.232 | 0.096 | 1.053 | 1.41 | 1.178 | 1.31 |
| extraction (spectral):classifier (NN) | 1.243 | 0.097 | 1.046 | 1.435 | 1.183 | 1.308 |
| extraction (LFCC):classifier (RF) | -0.878 | 0.104 | -1.07 | -0.663 | -0.929 | -0.819 |
| extraction (MFCC):classifier (RF) | -0.918 | 0.087 | -1.095 | -0.743 | -0.974 | -0.863 |
| extraction (spectral):classifier (RF) | -0.812 | 0.077 | -0.96 | -0.651 | -0.878 | -0.775 |
| extraction (LFCC):classifier (SVM) | 1.629 | 0.114 | 1.411 | 1.881 | 1.557 | 1.692 |
| extraction (MFCC):classifier (SVM) | 1.723 | 0.128 | 1.465 | 1.957 | 1.631 | 1.791 |
| extraction (spectral):classifier (SVM) | 1.908 | 0.125 | 1.668 | 2.156 | 1.827 | 1.981 |

Table S6b. Fitted values and upper and lower confidence limits on the response scale from the model with overall accuracy as the response for data including 16 different species with n=10 calls per individual, using only PCA data.

| Extraction method | Classifier | Estimate | Lower CI | Upper CI |
|---|---|---|---|---|
| HCTSA | DFA | 0.332 | 0.261 | 0.411 |
| HCTSA | NN | 0.11 | 0.072 | 0.163 |
| HCTSA | RF | 0.537 | 0.426 | 0.643 |
| HCTSA | SVM | 0.077 | 0.048 | 0.118 |
| LFCC | DFA | 0.632 | 0.406 | 0.814 |
| LFCC | NN | 0.587 | 0.29 | 0.834 |
| LFCC | RF | 0.624 | 0.33 | 0.853 |
| LFCC | SVM | 0.594 | 0.287 | 0.846 |
| MFCC | DFA | 0.659 | 0.482 | 0.802 |
| MFCC | NN | 0.622 | 0.37 | 0.822 |
| MFCC | RF | 0.643 | 0.395 | 0.832 |
| MFCC | SVM | 0.644 | 0.366 | 0.846 |
| spectral | DFA | 0.365 | 0.254 | 0.487 |
| spectral | NN | 0.331 | 0.177 | 0.526 |
| spectral | RF | 0.374 | 0.215 | 0.56 |
| spectral | SVM | 0.393 | 0.215 | 0.612 |

**Table S7. Model results (accuracy) for different sample sizes - n10 vs n30 vs n60.**

Table S7a. Results of the model with accuracy as the response (estimates, together with standard errors (SE), confidence limits (CI), as well as minimum and maximum of model estimates obtained when excluding individual call types one at a time) used to analyse the effects of sample size (4 different species with n=10, n=30 and n=60 calls per individual).

| Term | Estimate | SE | Lower CI | Upper CI | Min | Max |
|---|---|---|---|---|---|---|
| (Intercept) | -0.945 | 0.571 | -2.140 | 0.234 | -1.490 | -0.538 |
| extraction(LFCC) | 0.484 | 0.187 | 0.130 | 0.882 | 0.329 | 0.613 |
| extraction(MFCC) | 1.714 | 0.716 | 0.327 | 2.975 | 1.086 | 2.002 |
| extraction(spectral) | 0.292 | 0.095 | 0.103 | 0.492 | 0.137 | 0.605 |
| classifier(NN) | -1.234 | 0.096 | -1.412 | -1.059 | -1.517 | -0.695 |
| classifier(RF) | 1.175 | 0.125 | 0.934 | 1.411 | 1.082 | 1.274 |
| classifier(SVM) | 0.632 | 0.119 | 0.396 | 0.872 | 0.582 | 0.713 |
| n30 | 0.477 | 0.059 | 0.348 | 0.611 | 0.376 | 0.729 |
| n60 | -1.376 | 0.077 | -1.519 | -1.228 | -1.660 | -0.498 |
| extraction(LFCC):classifier(NN) | 0.979 | 0.093 | 0.790 | 1.150 | 0.394 | 1.294 |
| extraction(MFCC):classifier(NN) | 0.990 | 0.095 | 0.794 | 1.178 | 0.531 | 1.261 |
| extraction(spectral):classifier(NN) | 0.996 | 0.092 | 0.800 | 1.182 | 0.495 | 1.241 |
| extraction(LFCC):classifier(RF) | -0.942 | 0.083 | -1.110 | -0.773 | -1.073 | -0.860 |
| extraction(MFCC):classifier(RF) | -1.102 | 0.084 | -1.267 | -0.933 | -1.281 | -0.997 |
| extraction(spectral):classifier(RF) | -0.855 | 0.081 | -1.023 | -0.681 | -1.141 | -0.733 |
| extraction(LFCC):classifier(SVM) | -0.408 | 0.084 | -0.596 | -0.251 | -0.496 | -0.354 |
| extraction(MFCC):classifier(SVM) | -0.454 | 0.084 | -0.644 | -0.275 | -0.596 | -0.391 |
| extraction(spectral):classifier(SVM) | -0.437 | 0.081 | -0.620 | -0.269 | -0.682 | -0.323 |
| extraction(LFCC):n30 | 0.086 | 0.081 | -0.088 | 0.260 | -0.256 | 0.243 |
| extraction(MFCC):n30 | -0.074 | 0.084 | -0.259 | 0.095 | -0.296 | 0.075 |
| extraction(spectral):n30 | -0.304 | 0.081 | -0.467 | -0.128 | -0.546 | -0.202 |
| extraction(LFCC):n60 | 2.108 | 0.096 | 1.913 | 2.298 | 1.138 | 2.450 |
| extraction(MFCC):n60 | 1.902 | 0.099 | 1.704 | 2.093 | 1.077 | 2.188 |
| extraction(spectral):n60 | 1.554 | 0.095 | 1.364 | 1.747 | 0.710 | 1.823 |
| classifier(NN):n30 | -0.507 | 0.101 | -0.715 | -0.319 | -0.685 | -0.394 |
| classifier(RF):n30 | 0.167 | 0.081 | -0.002 | 0.330 | -0.141 | 0.276 |
| classifier(SVM):n30 | -0.135 | 0.080 | -0.310 | 0.037 | -0.421 | -0.003 |
| classifier(NN):n60 | 2.292 | 0.106 | 2.085 | 2.495 | 1.308 | 2.646 |
| classifier(RF):n60 | 2.132 | 0.096 | 1.943 | 2.324 | 1.215 | 2.473 |
| classifier(SVM):n60 | 1.580 | 0.095 | 1.388 | 1.765 | 0.705 | 1.892 |
| extraction(LFCC):classifier(NN):n30 | 0.566 | 0.129 | 0.309 | 0.829 | 0.448 | 0.821 |
| extraction(MFCC):classifier(NN):n30 | 0.592 | 0.133 | 0.325 | 0.872 | 0.496 | 0.787 |
| extraction(spectral):classifier(NN):n30 | 0.766 | 0.129 | 0.514 | 1.041 | 0.707 | 0.903 |
| extraction(LFCC):classifier(RF):n30 | -0.179 | 0.116 | -0.416 | 0.066 | -0.365 | 0.181 |
| extraction(MFCC):classifier(RF):n30 | -0.129 | 0.119 | -0.369 | 0.123 | -0.403 | 0.180 |
| extraction(spectral):classifier(RF):n30 | 0.003 | 0.114 | -0.250 | 0.239 | -0.092 | 0.266 |
| extraction(LFCC):classifier(SVM):n30 | 0.132 | 0.116 | -0.115 | 0.367 | 0.034 | 0.429 |
| extraction(MFCC):classifier(SVM):n30 | 0.215 | 0.118 | -0.032 | 0.453 | 0.003 | 0.468 |
| extraction(spectral):classifier(SVM):n30 | 0.414 | 0.114 | 0.174 | 0.663 | 0.251 | 0.649 |

| | | | | | | |
|---|---|---|---|---|---|---|
| extraction(LFCC):classifier(NN):n60 | -2.199 | 0.134 | -2.442 | -1.933 | -2.630 | -1.108 |
| extraction(MFCC):classifier(NN):n60 | -2.134 | 0.137 | -2.410 | -1.877 | -2.476 | -1.165 |
| extraction(spectral):classifier(NN):n60 | -1.967 | 0.134 | -2.237 | -1.719 | -2.290 | -1.040 |
| extraction(LFCC):classifier(RF):n60 | -2.209 | 0.127 | -2.459 | -1.933 | -2.615 | -1.244 |
| extraction(MFCC):classifier(RF):n60 | -1.996 | 0.130 | -2.277 | -1.718 | -2.425 | -1.087 |
| extraction(spectral):classifier(RF):n60 | -1.736 | 0.125 | -1.998 | -1.478 | -2.030 | -0.907 |
| extraction(LFCC):classifier(SVM):n60 | -1.588 | 0.127 | -1.845 | -1.334 | -1.885 | -0.718 |
| extraction(MFCC):classifier(SVM):n60 | -1.438 | 0.129 | -1.712 | -1.173 | -1.900 | -0.596 |
| extraction(spectral):classifier(SVM):n60 | -1.142 | 0.125 | -1.390 | -0.878 | -1.417 | -0.348 |

Table S7b. Fitted values from the model (backtransformed to the response scale) with accuracy as the response used to analyse the effects of sample size (4 different species with n=10, n=30 and n=60 calls per individual).

| Extraction method | Classifier | n | Accuracy | Lower CI | Upper CI |
|---|---|---|---|---|---|
| HCTSA | DFA | n10 | 0.280 | 0.105 | 0.558 |
| HCTSA | NN | n10 | 0.102 | 0.028 | 0.305 |
| HCTSA | RF | n10 | 0.557 | 0.23 | 0.838 |
| HCTSA | SVM | n10 | 0.422 | 0.149 | 0.751 |
| LFCC | DFA | n10 | 0.387 | 0.14 | 0.753 |
| LFCC | NN | n10 | 0.328 | 0.081 | 0.77 |
| LFCC | RF | n10 | 0.443 | 0.105 | 0.853 |
| LFCC | SVM | n10 | 0.441 | 0.113 | 0.85 |
| MFCC | DFA | n10 | 0.683 | 0.118 | 0.961 |
| MFCC | NN | n10 | 0.629 | 0.067 | 0.965 |
| MFCC | RF | n10 | 0.699 | 0.101 | 0.976 |
| MFCC | SVM | n10 | 0.721 | 0.099 | 0.978 |
| spectral | DFA | n10 | 0.342 | 0.115 | 0.674 |
| spectral | NN | n10 | 0.291 | 0.066 | 0.7 |
| spectral | RF | n10 | 0.418 | 0.107 | 0.811 |
| spectral | SVM | n10 | 0.388 | 0.094 | 0.791 |
| HCTSA | DFA | n30 | 0.385 | 0.143 | 0.7 |
| HCTSA | NN | n30 | 0.099 | 0.019 | 0.37 |
| HCTSA | RF | n30 | 0.706 | 0.297 | 0.93 |
| HCTSA | SVM | n30 | 0.507 | 0.154 | 0.852 |
| LFCC | DFA | n30 | 0.526 | 0.118 | 0.879 |
| LFCC | NN | n30 | 0.477 | 0.067 | 0.93 |
| LFCC | RF | n30 | 0.580 | 0.101 | 0.954 |
| LFCC | SVM | n30 | 0.580 | 0.099 | 0.953 |
| MFCC | DFA | n30 | 0.764 | 0.151 | 0.98 |
| MFCC | NN | n30 | 0.734 | 0.061 | 0.99 |
| MFCC | RF | n30 | 0.783 | 0.081 | 0.992 |
| MFCC | SVM | n30 | 0.807 | 0.09 | 0.993 |
| spectral | DFA | n30 | 0.382 | 0.104 | 0.77 |
| spectral | NN | n30 | 0.387 | 0.049 | 0.886 |

| | | | | | |
|---|---|---|---|---|---|
| spectral | RF | n30 | 0.503 | 0.076 | 0.925 |
| spectral | SVM | n30 | 0.499 | 0.075 | 0.925 |
| HCTSA | DFA | n60 | 0.089 | 0.025 | 0.27 |
| HCTSA | NN | n60 | 0.221 | 0.048 | 0.609 |
| HCTSA | RF | n60 | 0.728 | 0.314 | 0.939 |
| HCTSA | SVM | n60 | 0.473 | 0.133 | 0.838 |
| LFCC | DFA | n60 | 0.567 | 0.166 | 0.899 |
| LFCC | NN | n60 | 0.527 | 0.069 | 0.945 |
| LFCC | RF | n60 | 0.605 | 0.09 | 0.961 |
| LFCC | SVM | n60 | 0.619 | 0.093 | 0.962 |
| MFCC | DFA | n60 | 0.785 | 0.164 | 0.983 |
| MFCC | NN | n60 | 0.770 | 0.071 | 0.992 |
| MFCC | RF | n60 | 0.818 | 0.091 | 0.994 |
| MFCC | SVM | n60 | 0.834 | 0.1 | 0.995 |
| spectral | DFA | n60 | 0.384 | 0.1 | 0.776 |
| spectral | NN | n60 | 0.405 | 0.049 | 0.895 |
| spectral | RF | n60 | 0.560 | 0.088 | 0.944 |
| spectral | SVM | n60 | 0.540 | 0.082 | 0.939 |

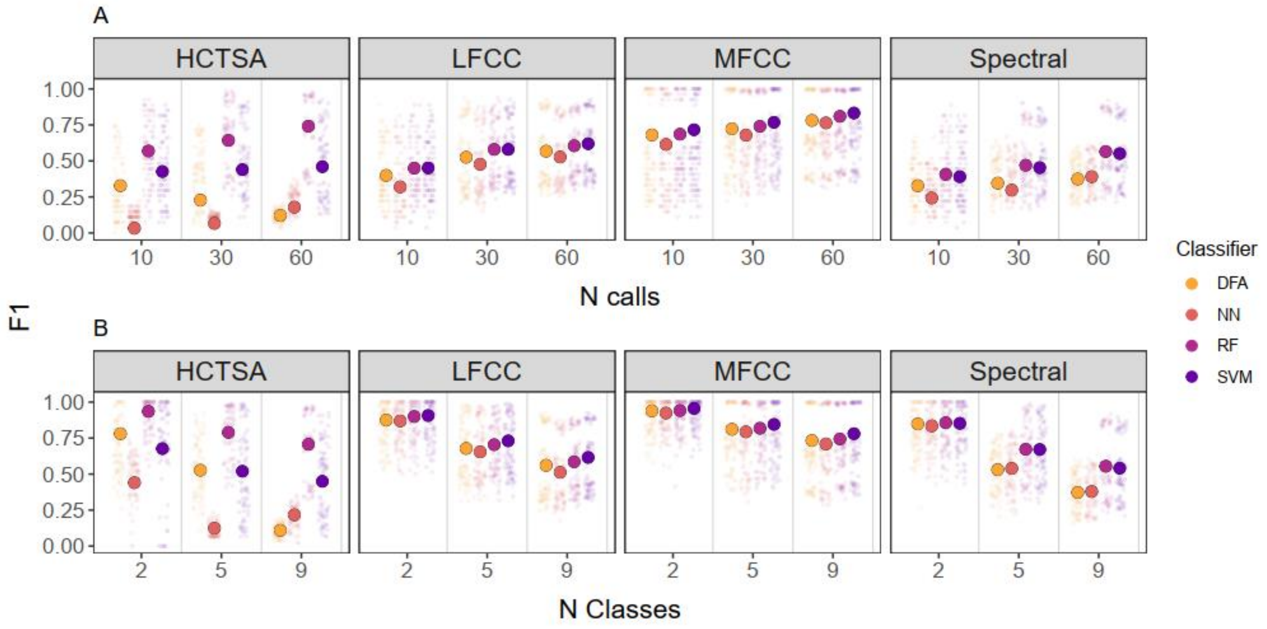


**Figure S4: F1 scores of machine learning algorithms classifying vocalisations to individual for datasets with different A) sample sizes (sample size of 10, 30, or 60 calls per individual, respectively); B) number of classes (2, 5 and 9 individuals per species).** Observed (smaller points) and model predicted estimates (larger points) for each classifier as plotted across various extraction methods. Dataset consists of vocalisations from 4 mammalian species. Values were back transformed to the response scale.

**Table S8. Model results (F1) for different sample sizes - n10 vs n30 vs n60.**

Table S8a. Results of the model with F1 as the response (estimates, together with standard errors (SE), confidence limits (Cl), as well as minimum and maximum of model estimates obtained when excluding individual call types one at a time) used to analyse the effects of sample size (4 different species with n=10, n=30 and n=60 calls per individual). Note that this model did not converge with the sample size predictor as a factor with three levels, so instead we included sample size as a z-transformed, numeric predictor.

| Term | Estimate | SE | Lower CI | Upper CI | Min | Max |
|---|---|---|---|---|---|---|
| (Intercept) | -1.305 | 0.563 | -2.362 | -0.265 | -1.396 | -0.890 |
| extraction(LFCC) | 1.239 | 0.185 | 0.881 | 1.571 | 0.895 | 1.384 |
| extraction(MFCC) | 2.301 | 0.667 | 1.028 | 3.535 | 1.331 | 2.599 |
| extraction(spectral) | 0.682 | 0.090 | 0.515 | 0.857 | 0.595 | 0.735 |
| classifier(NN) | -1.214 | 0.062 | -1.338 | -1.097 | -0.822 | -0.487 |
| classifier(RF) | 1.945 | 0.122 | 1.725 | 2.205 | 1.617 | 2.052 |
| classifier(SVM) | 1.072 | 0.117 | 0.852 | 1.317 | 0.796 | 1.232 |
| number of calls (z-transformed, numeric) | -0.518 | 0.033 | -0.591 | -0.446 | -0.616 | -0.214 |
| extraction(LFCC:)classifier(NN) | 0.941 | 0.068 | 0.813 | 1.079 | 0.299 | 0.603 |
| extraction(MFCC:)classifier(NN) | 1.016 | 0.072 | 0.889 | 1.150 | 0.314 | 0.659 |
| extraction(spectral):classifier(NN) | 1.023 | 0.068 | 0.891 | 1.160 | 0.448 | 0.758 |
| extraction(LFCC):classifier(RF) | -1.775 | 0.058 | -1.893 | -1.659 | -1.872 | -1.413 |
| extraction(MFCC):classifier(RF) | -1.850 | 0.061 | -1.969 | -1.734 | -1.944 | -1.567 |
| extraction(spectral):classifier(RF) | -1.400 | 0.057 | -1.520 | -1.286 | -1.453 | -1.341 |
| extraction(LFCC):classifier(SVM) | -0.859 | 0.058 | -0.974 | -0.740 | -1.047 | -0.582 |
| extraction(MFCC):classifier(SVM) | -0.825 | 0.061 | -0.950 | -0.702 | -1.017 | -0.627 |
| extraction(spectral):classifier(SVM) | -0.591 | 0.057 | -0.709 | -0.477 | -0.725 | -0.572 |
| extraction(LFCC):(z)ncalls | 0.820 | 0.041 | 0.734 | 0.906 | 0.460 | 0.931 |
| extraction(MFCC):(z).ncalls | 0.731 | 0.043 | 0.641 | 0.821 | 0.441 | 0.814 |
| extraction(spectral):(z)ncalls | 0.602 | 0.041 | 0.512 | 0.688 | 0.295 | 0.677 |
| classifier(NN):z.n | 1.283 | 0.054 | 1.184 | 1.384 | 0.575 | 1.084 |
| classifier(RF):z.n | 0.835 | 0.042 | 0.749 | 0.920 | 0.493 | 0.952 |
| classifier(SVM):z.n | 0.573 | 0.041 | 0.492 | 0.660 | 0.286 | 0.693 |
| extraction(LFCC):classifier(NN):(z)ncalls | -1.218 | 0.066 | -1.345 | -1.083 | -1.076 | -0.500 |
| extraction(MFCC):classifier(NN):(z)ncalls | -1.205 | 0.068 | -1.345 | -1.071 | -1.016 | -0.520 |
| extraction(spectral):classifier(NN):(z)ncalls | -1.082 | 0.067 | -1.206 | -0.960 | -0.949 | -0.475 |
| extraction(LFCC):classifier(RF):(z)ncalls | -0.868 | 0.057 | -0.985 | -0.745 | -1.010 | -0.507 |
| extraction(MFCC):classifier(RF):(z)ncalls | -0.775 | 0.059 | -0.893 | -0.650 | -0.898 | -0.441 |
| extraction(spectral):classifier(RF):(z)ncalls | -0.657 | 0.056 | -0.769 | -0.533 | -0.773 | -0.368 |
| extraction(LFCC):classifier(SVM):(z)ncalls | -0.575 | 0.057 | -0.689 | -0.463 | -0.690 | -0.291 |
| extraction(MFCC):classifier(SVM):(z)ncalls | -0.509 | 0.058 | -0.637 | -0.391 | -0.691 | -0.242 |
| extraction(spectral):classifier(SVM):(z)ncalls | -0.391 | 0.056 | -0.515 | -0.267 | -0.508 | -0.146 |

Table S8b. Fitted values from the model (backtransformed to the response scale) with accuracy as the response used to analyse the effects of sample size (n; 4 different species with n=10, n=30 and n=60 calls per individual).

| Extraction method | Classifier | n | F1 | Lower CI | Upper CI |
|---|---|---|---|---|---|
| HCTSA | DFA | n10 | 0.328 | 0.156 | 0.56 |
| HCTSA | NN | n10 | 0.033 | 0.012 | 0.081 |
| HCTSA | RF | n10 | 0.570 | 0.306 | 0.802 |
| HCTSA | SVM | n10 | 0.426 | 0.198 | 0.692 |
| LFCC | DFA | n10 | 0.399 | 0.199 | 0.753 |
| LFCC | NN | n10 | 0.320 | 0.16 | 0.77 |
| LFCC | RF | n10 | 0.450 | 0.187 | 0.853 |
| LFCC | SVM | n10 | 0.452 | 0.21 | 0.85 |
| MFCC | DFA | n10 | 0.680 | 0.118 | 0.945 |
| MFCC | NN | n10 | 0.615 | 0.067 | 0.927 |
| MFCC | RF | n10 | 0.686 | 0.101 | 0.953 |
| MFCC | SVM | n10 | 0.717 | 0.099 | 0.959 |
| spectral | DFA | n10 | 0.328 | 0.147 | 0.579 |
| spectral | NN | n10 | 0.243 | 0.102 | 0.475 |
| spectral | RF | n10 | 0.407 | 0.178 | 0.689 |
| spectral | SVM | n10 | 0.391 | 0.17 | 0.671 |
| HCTSA | DFA | n30 | 0.228 | 0.094 | 0.452 |
| HCTSA | NN | n30 | 0.066 | 0.022 | 0.18 |
| HCTSA | RF | n30 | 0.643 | 0.34 | 0.866 |
| HCTSA | SVM | n30 | 0.440 | 0.183 | 0.735 |
| LFCC | DFA | n30 | 0.526 | 0.118 | 0.879 |
| LFCC | NN | n30 | 0.477 | 0.067 | 0.93 |
| LFCC | RF | n30 | 0.580 | 0.101 | 0.954 |
| LFCC | SVM | n30 | 0.580 | 0.099 | 0.953 |
| MFCC | DFA | n30 | 0.724 | 0.207 | 0.961 |
| MFCC | NN | n30 | 0.679 | 0.146 | 0.961 |
| MFCC | RF | n30 | 0.740 | 0.173 | 0.974 |
| MFCC | SVM | n30 | 0.768 | 0.195 | 0.978 |
| spectral | DFA | n30 | 0.346 | 0.138 | 0.635 |
| spectral | NN | n30 | 0.297 | 0.093 | 0.634 |
| spectral | RF | n30 | 0.470 | 0.164 | 0.804 |
| spectral | SVM | n30 | 0.454 | 0.156 | 0.791 |
| HCTSA | DFA | n60 | 0.122 | 0.042 | 0.301 |
| HCTSA | NN | n60 | 0.179 | 0.051 | 0.464 |
| HCTSA | RF | n60 | 0.741 | 0.394 | 0.928 |
| HCTSA | SVM | n60 | 0.460 | 0.163 | 0.791 |
| LFCC | DFA | n60 | 0.567 | 0.166 | 0.899 |
| LFCC | NN | n60 | 0.527 | 0.069 | 0.945 |
| LFCC | RF | n60 | 0.605 | 0.09 | 0.961 |
| LFCC | SVM | n60 | 0.619 | 0.093 | 0.962 |
| MFCC | DFA | n60 | 0.781 | 0.219 | 0.977 |
| MFCC | NN | n60 | 0.764 | 0.127 | 0.985 |
| MFCC | RF | n60 | 0.810 | 0.154 | 0.99 |

| | | | | | |
|---|---|---|---|---|---|
| MFCC | SVM | n60 | 0.833 | 0.174 | 0.991 |
| spectral | DFA | n60 | 0.374 | 0.125 | 0.712 |
| spectral | NN | n60 | 0.391 | 0.081 | 0.821 |
| spectral | RF | n60 | 0.565 | 0.145 | 0.911 |
| spectral | SVM | n60 | 0.551 | 0.137 | 0.905 |

**Table S9. Model results (accuracy) for different number of individuals – 2 vs 5 vs 9 classes.**

Table S9a. Results of the model with accuracy as the response (estimates, together with standard errors (SE), confidence limits (Cl), as well as minimum and maximum of model estimates obtained when excluding individual call types one at a time) used to analyse the effects of different number of individuals (4 different species with 2, 5, or 9 individuals per species).

| Term | Estimate | SE | Lower CI | Upper CI | Min | Max |
|---|---|---|---|---|---|---|
| (Intercept) | 1.321 | 0.549 | 0.139 | 2.101 | -1.490 | -0.538 |
| extraction(LFCC) | 0.730 | 0.250 | 0.253 | 1.257 | 0.329 | 0.613 |
| extraction(MFCC) | 1.605 | 0.526 | 0.484 | 2.490 | 1.086 | 2.002 |
| extraction(spectral) | 0.480 | 0.140 | 0.191 | 0.748 | 0.137 | 0.605 |
| classifier(NN) | -1.283 | 0.112 | -1.505 | -1.054 | -1.517 | -0.695 |
| classifier(RF) | 1.488 | 0.150 | 1.199 | 1.793 | 1.082 | 1.274 |
| classifier(SVM) | 0.170 | 0.166 | -0.145 | 0.489 | 0.582 | 0.713 |
| classes(5individuals) | -1.174 | 0.167 | -1.504 | -0.848 | 0.376 | 0.729 |
| classes(9individuals) | -3.383 | 0.190 | -3.772 | -2.991 | -1.660 | -0.498 |
| extraction(LFCC:)classifier(NN) | 1.207 | 0.128 | 0.961 | 1.463 | 0.394 | 1.294 |
| extraction(MFCC:)classifier(NN) | 1.001 | 0.137 | 0.732 | 1.306 | 0.531 | 1.261 |
| extraction(spectral):classifier(NN) | 1.155 | 0.124 | 0.917 | 1.416 | 0.495 | 1.241 |
| extraction(LFCC):classifier(RF) | -1.185 | 0.144 | -1.464 | -0.884 | -1.073 | -0.860 |
| extraction(MFCC):classifier(RF) | -1.414 | 0.151 | -1.716 | -1.089 | -1.281 | -0.997 |
| extraction(spectral):classifier(RF) | -1.401 | 0.139 | -1.675 | -1.123 | -1.141 | -0.733 |
| extraction(LFCC):classifier(SVM) | 0.203 | 0.132 | -0.058 | 0.492 | -0.496 | -0.354 |
| extraction(MFCC):classifier(SVM) | 0.168 | 0.142 | 0.139 | 2.101 | -0.596 | -0.391 |
| extraction(spectral):classifier(SVM) | -0.141 | 0.127 | 0.253 | 1.257 | -0.682 | -0.323 |
| extraction(LFCC):classes(5individuals) | -0.081 | 0.120 | 0.484 | 2.490 | -0.256 | 0.243 |
| extraction(MFCC):classes(5individuals) | -0.207 | 0.128 | 0.191 | 0.748 | -0.296 | 0.075 |
| extraction(spectral):classes(5individuals) | -0.444 | 0.118 | -1.505 | -1.054 | -0.546 | -0.202 |
| extraction(LFCC):classes(9individuals) | 1.600 | 0.133 | 1.199 | 1.793 | 1.138 | 2.450 |
| extraction(MFCC):classes(9individuals) | 1.525 | 0.140 | -0.145 | 0.489 | 1.077 | 2.188 |
| extraction(spectral):classes(9individuals) | 1.121 | 0.132 | -1.504 | -0.848 | 0.710 | 1.823 |
| classifier(NN):classes(5individuals) | -0.262 | 0.119 | -3.772 | -2.991 | -0.685 | -0.394 |
| classifier(RF):classes(5individuals) | -0.254 | 0.131 | 0.961 | 1.463 | -0.141 | 0.276 |
| classifier(SVM):classes(5individuals) | -0.017 | 0.115 | 0.732 | 1.306 | -0.421 | -0.003 |
| classifier(NN):classes(5individuals) | 2.178 | 0.130 | 0.917 | 1.416 | 1.308 | 2.646 |
| classifier(RF):classes(5individuals) | 1.506 | 0.141 | -1.464 | -0.884 | 1.215 | 2.473 |
| classifier(SVM):classes(5individuals) | 1.799 | 0.129 | -1.716 | -1.089 | 0.705 | 1.892 |
| extraction(LFCC):classifier(NN):classes(5individuals) | 0.225 | 0.173 | -1.675 | -1.123 | 0.448 | 0.821 |
| extraction(MFCC):classifier(NN):classes(5individuals) | 0.431 | 0.182 | -0.058 | 0.492 | 0.496 | 0.787 |
| extraction(spectral):classifier(NN):classes(5individuals) | 0.448 | 0.170 | 0.139 | 2.101 | 0.707 | 0.903 |
| extraction(LFCC):classifier(RF):classes(5individuals) | 0.107 | 0.185 | 0.253 | 1.257 | -0.365 | 0.181 |
| extraction(MFCC):classifier(RF):classes(5individuals) | 0.287 | 0.193 | 0.484 | 2.490 | -0.403 | 0.180 |
| extraction(spectral):classifier(RF):classes(5individuals) | 0.747 | 0.180 | 0.191 | 0.748 | -0.092 | 0.266 |
| extraction(LFCC):classifier(SVM):classes(5individuals) | -0.094 | 0.174 | -1.505 | -1.054 | 0.034 | 0.429 |
| extraction(MFCC):classifier(SVM):classes(5individuals) | -0.055 | 0.183 | 1.199 | 1.793 | 0.003 | 0.468 |
| extraction(spectral):classifier(SVM):classes(5individuals) | 0.553 | 0.168 | -0.145 | 0.489 | 0.251 | 0.649 |

| | | | | | | |
|---|---|---|---|---|---|---|
| extraction(LFCC):classifier(NN):classes(9individuals) | -2.262 | 0.179 | -1.504 | -0.848 | -2.630 | -1.108 |
| extraction(MFCC):classifier(NN):classes(9individuals) | -1.977 | 0.187 | -3.772 | -2.991 | -2.476 | -1.165 |
| extraction(spectral):classifier(NN):classes(9individuals) | -1.964 | 0.177 | 0.961 | 1.463 | -2.290 | -1.040 |
| extraction(LFCC):classifier(RF):classes(9individuals) | -1.659 | 0.191 | 0.732 | 1.306 | -2.615 | -1.244 |
| extraction(MFCC):classifier(RF):classes(9individuals) | -1.460 | 0.198 | 0.917 | 1.416 | -2.425 | -1.087 |
| extraction(spectral):classifier(RF):classes(9individuals) | -0.889 | 0.187 | -1.464 | -0.884 | -2.030 | -0.907 |
| extraction(LFCC):classifier(SVM):classes(9individuals) | -1.932 | 0.182 | -1.716 | -1.089 | -1.885 | -0.718 |
| extraction(MFCC):classifier(SVM):classes(9individuals) | -1.854 | 0.190 | -1.675 | -1.123 | -1.900 | -0.596 |
| extraction(spectral):classifier(SVM):classes(9individuals) | -1.190 | 0.177 | -0.058 | 0.492 | -1.417 | -0.348 |

Table S9b. Fitted values from the model (backtransformed to the response scale) with accuracy as the response used to analyse the effects of different number of individuals (4 different species with 2, 5, or 9 individuals per species).

| Extraction method | Classifier | n individuals | Accuracy | Lower CI | Upper CI |
|---|---|---|---|---|---|
| HCTSA | DFA | 2cl | 0.789 | 0.535 | 0.891 |
| HCTSA | NN | 2cl | 0.509 | 0.203 | 0.74 |
| HCTSA | RF | 2cl | 0.943 | 0.792 | 0.98 |
| HCTSA | SVM | 2cl | 0.816 | 0.499 | 0.93 |
| LFCC | DFA | 2cl | 0.886 | 0.597 | 0.966 |
| LFCC | NN | 2cl | 0.878 | 0.462 | 0.977 |
| LFCC | RF | 2cl | 0.913 | 0.532 | 0.986 |
| LFCC | SVM | 2cl | 0.919 | 0.547 | 0.987 |
| MFCC | DFA | 2cl | 0.949 | 0.651 | 0.99 |
| MFCC | NN | 2cl | 0.934 | 0.463 | 0.992 |
| MFCC | RF | 2cl | 0.953 | 0.527 | 0.995 |
| MFCC | SVM | 2cl | 0.963 | 0.587 | 0.996 |
| spectral | DFA | 2cl | 0.858 | 0.582 | 0.945 |
| spectral | NN | 2cl | 0.842 | 0.436 | 0.961 |
| spectral | RF | 2cl | 0.868 | 0.464 | 0.971 |
| spectral | SVM | 2cl | 0.862 | 0.451 | 0.969 |
| HCTSA | DFA | 5cl | 0.537 | 0.203 | 0.778 |
| HCTSA | NN | 5cl | 0.198 | 0.034 | 0.546 |
| HCTSA | RF | 5cl | 0.799 | 0.337 | 0.955 |
| HCTSA | SVM | 5cl | 0.574 | 0.148 | 0.879 |
| LFCC | DFA | 5cl | 0.689 | 0.205 | 0.978 |
| LFCC | NN | 5cl | 0.664 | 0.07 | 0.992 |
| LFCC | RF | 5cl | 0.721 | 0.075 | 0.995 |
| LFCC | SVM | 5cl | 0.742 | 0.091 | 0.995 |
| MFCC | DFA | 5cl | 0.824 | 0.194 | 0.936 |
| MFCC | NN | 5cl | 0.807 | 0.071 | 0.975 |
| MFCC | RF | 5cl | 0.839 | 0.076 | 0.983 |
| MFCC | SVM | 5cl | 0.859 | 0.088 | 0.984 |
| spectral | DFA | 5cl | 0.545 | 0.134 | 0.86 |
| spectral | NN | 5cl | 0.560 | 0.055 | 0.951 |
| spectral | RF | 5cl | 0.682 | 0.077 | 0.974 |

| | | | | | |
|---|---|---|---|---|---|
| spectral | SVM | 5cl | 0.678 | 0.079 | 0.972 |
| HCTSA | DFA | 9cl | 0.113 | 0.026 | 0.291 |
| HCTSA | NN | 9cl | 0.237 | 0.039 | 0.621 |
| HCTSA | RF | 9cl | 0.717 | 0.23 | 0.937 |
| HCTSA | SVM | 9cl | 0.477 | 0.097 | 0.841 |
| LFCC | DFA | 9cl | 0.567 | 0.131 | 0.968 |
| LFCC | NN | 9cl | 0.527 | 0.043 | 0.989 |
| LFCC | RF | 9cl | 0.603 | 0.045 | 0.992 |
| LFCC | SVM | 9cl | 0.624 | 0.054 | 0.993 |
| MFCC | DFA | 9cl | 0.744 | 0.115 | 0.905 |
| MFCC | NN | 9cl | 0.728 | 0.036 | 0.961 |
| MFCC | RF | 9cl | 0.767 | 0.042 | 0.975 |
| MFCC | SVM | 9cl | 0.794 | 0.047 | 0.977 |
| spectral | DFA | 9cl | 0.387 | 0.07 | 0.776 |
| spectral | NN | 9cl | 0.407 | 0.027 | 0.919 |
| spectral | RF | 9cl | 0.560 | 0.043 | 0.962 |
| spectral | SVM | 9cl | 0.544 | 0.042 | 0.956 |

**Table S10. Model results (F1) for different number of individuals – 2 vs 5 vs 9 classes.**

Table S10. Classes F1 Results of the model with F1 as the response (estimates, together with standard errors (SE), confidence limits (Cl), as well as minimum and maximum of model estimates obtained when excluding individual call types one at a time) used to analyse the effects of different number of individuals (4 different species with 2, 5, or 9 individuals per species).

| Term | Estimate | SE | Lower Cl | Upper Cl | Min | Max |
|---|---|---|---|---|---|---|
| (Intercept) | 1.268 | 0.520 | 0.107 | 1.916 | 0.814 | 1.587 |
| extraction(LFCC) | 0.680 | 0.233 | 0.248 | 1.131 | 0.541 | 0.787 |
| extraction(MFCC) | 1.469 | 0.458 | 0.576 | 2.224 | 1.183 | 1.634 |
| extraction(spectral) | 0.454 | 0.144 | 0.152 | 0.758 | 0.336 | 0.596 |
| classifier(NN) | -1.507 | 0.114 | -1.727 | -1.290 | -1.876 | -1.145 |
| classifier(RF) | 1.402 | 0.149 | 1.107 | 1.703 | 1.244 | 1.525 |
| classifier(SVM) | -0.529 | 0.181 | -0.857 | -0.201 | -0.882 | -0.049 |
| classes(5individuals) | -1.161 | 0.173 | -1.497 | -0.801 | -1.319 | -0.953 |
| classes(9individuals) | -3.368 | 0.191 | -3.732 | -2.996 | -3.747 | -2.603 |
| extraction(LFCC:)classifier(NN) | 1.449 | 0.149 | 1.175 | 1.713 | 1.144 | 1.792 |
| extraction(MFCC:)classifier(NN) | 1.266 | 0.158 | 0.952 | 1.579 | 0.922 | 1.768 |
| extraction(spectral):classifier(NN) | 1.402 | 0.146 | 1.102 | 1.695 | 1.049 | 1.749 |
| extraction(LFCC):classifier(RF) | -1.153 | 0.163 | -1.463 | -0.837 | -1.386 | -1.004 |
| extraction(MFCC):classifier(RF) | -1.364 | 0.170 | -1.718 | -1.042 | -1.549 | -1.190 |
| extraction(spectral):classifier(RF) | -1.334 | 0.159 | -1.639 | -1.002 | -1.430 | -1.193 |
| extraction(LFCC):classifier(SVM) | 0.855 | 0.149 | 0.554 | 1.154 | 0.371 | 1.152 |
| extraction(MFCC):classifier(SVM) | 0.867 | 0.159 | 0.544 | 1.199 | 0.436 | 1.135 |
| extraction(spectral):classifier(SVM) | 0.557 | 0.144 | 0.251 | 0.847 | 0.132 | 0.851 |
| extraction(LFCC):classes(5individuals) | -0.041 | 0.141 | -0.325 | 0.237 | -0.141 | 0.128 |
| extraction(MFCC):classes(5individuals) | -0.119 | 0.149 | -0.425 | 0.217 | -0.418 | 0.110 |
| extraction(spectral):classes(5individuals) | -0.439 | 0.140 | -0.749 | -0.143 | -0.606 | -0.297 |
| extraction(LFCC):classes(9individuals) | 1.661 | 0.156 | 1.344 | 1.959 | 1.003 | 2.056 |
| extraction(MFCC):classes(9individuals) | 1.641 | 0.163 | 1.354 | 1.963 | 0.760 | 2.097 |
| extraction(spectral):classes(9individuals) | 1.124 | 0.156 | 0.831 | 1.438 | 0.730 | 1.370 |
| classifier(NN):classes(5individuals) | -0.555 | 0.147 | -0.861 | -0.261 | -0.735 | -0.347 |
| classifier(RF):classes(5individuals) | -0.188 | 0.151 | -0.517 | 0.131 | -0.342 | -0.098 |
| classifier(SVM):classes(5individuals) | 0.501 | 0.131 | 0.243 | 0.762 | 0.203 | 0.720 |
| classifier(NN):classes(5individuals) | 2.320 | 0.155 | 2.042 | 2.623 | 1.492 | 2.801 |
| classifier(RF):classes(5individuals) | 1.578 | 0.163 | 1.242 | 1.893 | 0.838 | 1.875 |
| classifier(SVM):classes(5individuals) | 2.427 | 0.149 | 2.152 | 2.722 | 2.039 | 2.859 |
| extraction(LFCC):classifier(NN):classes(5individuals) | 0.504 | 0.208 | 0.090 | 0.930 | 0.343 | 0.677 |
| extraction(MFCC):classifier(NN):classes(5individuals) | 0.685 | 0.216 | 0.241 | 1.130 | 0.322 | 0.825 |
| extraction(spectral):classifier(NN):classes(5individuals) | 0.695 | 0.205 | 0.286 | 1.123 | 0.514 | 0.838 |
| extraction(LFCC):classifier(RF):classes(5individuals) | 0.061 | 0.214 | -0.358 | 0.481 | -0.038 | 0.206 |
| extraction(MFCC):classifier(RF):classes(5individuals) | 0.198 | 0.221 | -0.297 | 0.668 | 0.050 | 0.464 |
| extraction(spectral):classifier(RF):classes(5individuals) | 0.715 | 0.209 | 0.285 | 1.144 | 0.646 | 0.802 |
| extraction(LFCC):classifier(SVM):classes(5individuals) | -0.576 | 0.200 | -1.011 | -0.177 | -0.726 | -0.271 |
| extraction(MFCC):classifier(SVM):classes(5individuals) | -0.604 | 0.210 | -1.052 | -0.145 | -0.864 | -0.381 |
| extraction(spectral):classifier(SVM):classes(5individuals) | 0.059 | 0.195 | -0.341 | 0.491 | -0.211 | 0.385 |

| | | | | | | |
|---|---|---|---|---|---|---|
| extraction(LFCC):classifier(NN):classes(9individuals) | -2.455 | 0.212 | -2.861 | -2.044 | -2.934 | -1.642 |
| extraction(MFCC):classifier(NN):classes(9individuals) | -2.197 | 0.219 | -2.636 | -1.785 | -2.800 | -1.343 |
| extraction(spectral):classifier(NN):classes(9individuals) | -2.187 | 0.211 | -2.592 | -1.779 | -2.656 | -1.389 |
| extraction(LFCC):classifier(RF):classes(9individuals) | -1.724 | 0.221 | -2.140 | -1.276 | -2.023 | -0.970 |
| extraction(MFCC):classifier(RF):classes(9individuals) | -1.561 | 0.227 | -1.999 | -1.101 | -1.931 | -0.693 |
| extraction(spectral):classifier(RF):classes(9individuals) | -0.906 | 0.217 | -1.336 | -0.468 | -1.140 | -0.394 |
| extraction(LFCC):classifier(SVM):classes(9individuals) | -2.525 | 0.211 | -2.937 | -2.131 | -2.927 | -2.112 |
| extraction(MFCC):classifier(SVM):classes(9individuals) | -2.508 | 0.219 | -2.982 | -2.063 | -2.970 | -2.037 |
| extraction(spectral):classifier(SVM):classes(9individuals) | -1.770 | 0.207 | -2.194 | -1.365 | -2.143 | -1.563 |

Table S10b. Fitted values from the model (back-transformed to the response scale) with F1 as the response used to analyse the effects of different number of individuals (4 different species with 2, 5, or 9 individuals per species).

| Extraction method | Classifier | n individuals | F1 | Lower CI | Upper CI |
|---|---|---|---|---|---|
| HCTSA | DFA | 2cl | 0.780 | 0.527 | 0.872 |
| HCTSA | NN | 2cl | 0.441 | 0.165 | 0.651 |
| HCTSA | RF | 2cl | 0.935 | 0.771 | 0.974 |
| HCTSA | SVM | 2cl | 0.677 | 0.321 | 0.847 |
| LFCC | DFA | 2cl | 0.875 | 0.588 | 0.955 |
| LFCC | NN | 2cl | 0.869 | 0.451 | 0.97 |
| LFCC | RF | 2cl | 0.900 | 0.5 | 0.98 |
| LFCC | SVM | 2cl | 0.907 | 0.513 | 0.982 |
| MFCC | DFA | 2cl | 0.939 | 0.664 | 0.984 |
| MFCC | NN | 2cl | 0.924 | 0.477 | 0.988 |
| MFCC | RF | 2cl | 0.941 | 0.518 | 0.992 |
| MFCC | SVM | 2cl | 0.956 | 0.591 | 0.994 |
| spectral | DFA | 2cl | 0.848 | 0.564 | 0.935 |
| spectral | NN | 2cl | 0.834 | 0.409 | 0.956 |
| spectral | RF | 2cl | 0.857 | 0.432 | 0.967 |
| spectral | SVM | 2cl | 0.852 | 0.414 | 0.965 |
| HCTSA | DFA | 5cl | 0.527 | 0.199 | 0.753 |
| HCTSA | NN | 5cl | 0.124 | 0.018 | 0.393 |
| HCTSA | RF | 5cl | 0.789 | 0.31 | 0.95 |
| HCTSA | SVM | 5cl | 0.520 | 0.119 | 0.842 |
| LFCC | DFA | 5cl | 0.678 | 0.225 | 0.972 |
| LFCC | NN | 5cl | 0.654 | 0.067 | 0.991 |
| LFCC | RF | 5cl | 0.704 | 0.065 | 0.993 |
| LFCC | SVM | 5cl | 0.731 | 0.086 | 0.994 |
| MFCC | DFA | 5cl | 0.811 | 0.187 | 0.923 |
| MFCC | NN | 5cl | 0.794 | 0.058 | 0.973 |
| MFCC | RF | 5cl | 0.818 | 0.063 | 0.981 |
| MFCC | SVM | 5cl | 0.845 | 0.073 | 0.982 |
| spectral | DFA | 5cl | 0.531 | 0.121 | 0.849 |
| spectral | NN | 5cl | 0.540 | 0.04 | 0.952 |
| spectral | RF | 5cl | 0.672 | 0.06 | 0.976 |

| | | | | | |
|---|---|---|---|---|---|
| spectral | SVM | 5cl | 0.671 | 0.063 | 0.974 |
| HCTSA | DFA | 9cl | 0.109 | 0.026 | 0.254 |
| HCTSA | NN | 9cl | 0.216 | 0.035 | 0.563 |
| HCTSA | RF | 9cl | 0.707 | 0.218 | 0.925 |
| HCTSA | SVM | 9cl | 0.450 | 0.089 | 0.809 |
| LFCC | DFA | 9cl | 0.560 | 0.155 | 0.957 |
| LFCC | NN | 9cl | 0.512 | 0.045 | 0.986 |
| LFCC | RF | 9cl | 0.585 | 0.045 | 0.99 |
| LFCC | SVM | 9cl | 0.615 | 0.055 | 0.992 |
| MFCC | DFA | 9cl | 0.733 | 0.116 | 0.882 |
| MFCC | NN | 9cl | 0.709 | 0.032 | 0.953 |
| MFCC | RF | 9cl | 0.744 | 0.036 | 0.971 |
| MFCC | SVM | 9cl | 0.780 | 0.042 | 0.972 |
| spectral | DFA | 9cl | 0.373 | 0.067 | 0.753 |
| spectral | NN | 9cl | 0.379 | 0.022 | 0.914 |
| spectral | RF | 9cl | 0.554 | 0.037 | 0.962 |
| spectral | SVM | 9cl | 0.541 | 0.036 | 0.958 |
